# Targeting protein glycosylation and cholesterol metabolism in chemoresistant pancreatic cancer

**DOI:** 10.1101/2025.04.13.648567

**Authors:** Yunguang Li, Shijie Tang, Huan Wang, Hongwen Zhu, Shiwei Guo, Yehan Zhang, Yurun Lu, Maoyuan Sun, Juan He, Yikai Li, Yi Zhang, Xiaohan Shi, Yuanxiang Miao, Chaoliang Zhong, Yiqin Zhu, Yi Ju, Yuejia Liu, Yong Wang, Luonan Chen, Hu Zhou, Gang Jin, Dong Gao

**Author notes:** Corresponding author. (Y.W.), (L.N.C.), (H.Z.), (G.J.), (D.G.). These authors contributed equally (Yunguang Li, Shijie Tang, Huan Wang, Hongwen Zhu, Shiwei Guo, Yehan Zhang, Yurun Lu).

## Abstract

Chemotherapy remains the cornerstone of pancreatic ductal adenocarcinoma (PDAC) treatment. However, most cases of PDAC finally progress to advanced chemoresistant disease. This highlights an urgent need to develop new pharmacological strategies to overcome chemotherapy resistance using clinical grade inhibitors. Here, we established a biobank comprising 260 organoid lines derived from 322 pancreatic cancer patients. These organoids underwent extensive multi-omics profiling and drug sensitivity testing for both chemotherapy and targeted therapy. Increased protein glycosylation and cholesterol metabolism signaling pathways were especially enriched in chemoresistant organoid lines. Importantly, chemoresistant PDAC organoids were effectively targeted with statins, inhibitors of cholesterol synthesis. Statins treatment attenuated protein glycosylation, cholesterol levels, and the epithelial-to-mesenchymal transition (EMT) signature in PDAC organoids. We further conducted a phase 2 clinical trial (NCT06241352) combining atorvastatin with chemotherapy in patients with advanced chemoresistant pancreatic cancer. Among 29 patients, 21 (72.4%) demonstrated a response, with tumor markers decreasing by more than 20%, suggesting durable responses and potential clinical benefits in this challenging patient population. ClinicalTrials.gov identifier: NCT06241352.

## Introduction

PDAC is the third leading cause of cancer-related death^1^. The primary treatment for PDAC remains cytotoxic chemotherapy. Current standard adjuvant treatments following surgical resection of PDAC include single-agent gemcitabine (GEM), 5-fluorouracil (5-FU), combinations such as gemcitabine/capecitabine, nab-paclitaxel/gemcitabine (AG) and FOLFIRINOX (FFX)^2–4^. Despite these aggressive treatment regimens, recurrence rates are notably high due to poor chemotherapy response rates and the development of therapeutic resistance, which is a major contributor to PDAC-related mortality^1^. Therefore, the establishment of preclinical models that accurately reflect the molecular characteristics of patient tumors is crucial for investigating the mechanisms underlying intrinsic and acquired chemotherapy resistance. Furthermore, there is an urgent need to develop novel pharmacological strategies to enhance and prolong chemotherapy responses in clinical settings.

Patient-derived cancer organoids provide a promising avenue for preclinical studies to characterize tumor behavior and investigate the molecular mechanisms of drug resistance^5,6^. Recently, several research groups, including ours, have generated patient-derived organoid biobanks from PDAC patients^7–9^. These organoids faithfully replicate the therapeutic responses observed in patients, supporting their potential for clinical translation^10,11^. Here, we generated 260 organoid lines from 322 pancreatic cancer patients to identify molecular signatures associated with chemotherapy response and to develop therapeutic strategies with potential clinical application. We found that, in chemoresistant organoid lines, protein glycosylation and cholesterol metabolism signaling activity were especially increased. Using high-throughput drug screening, we identified statins, inhibitors of HMG CoA reductase (HMGCR) commonly used to lower cholesterol levels, as effective agents for enhancing chemotherapy response in chemoresistant PDAC organoids. In a phase 2 clinical trial combining atorvastatin with chemotherapy, 72.4% of patients with advanced chemoresistant pancreatic cancer demonstrated a response, with tumor markers decreasing by more than 20%.

## Results

### Proteogenomic profiling of PDAC

Pancreatic cancer samples were collected through surgical resection, endoscopic ultrasound-guided fine-needle aspiration biopsy (EUS-FNA), or ascites collection, leading to the establishment of 260 organoid lines derived from 322 patients (Fig. 1a). These organoid lines originated from various tumor types, including 239 PDACs, 10 adenosquamous carcinomas, 7 intraductal papillary mucinous neoplasms, 3 acinar cell carcinomas, and 1 solid pseudopapillary tumor (Supplementary Table 1). The patient cohort includes both early-stage (Stage I/II, ∼70%) and advanced-stage (Stage III/IV, ∼30%) pancreatic cancers (Supplementary Table 1). The demographic distribution of this patient cohort was consistent with previous large-scale studies, with 60% male patients and 57% of the samples originating from the pancreatic head region (Supplementary Table 1). Clinical and pathological details of the patients corresponding to the organoids are summarized in Supplementary Table 1.

**Fig. 1:**
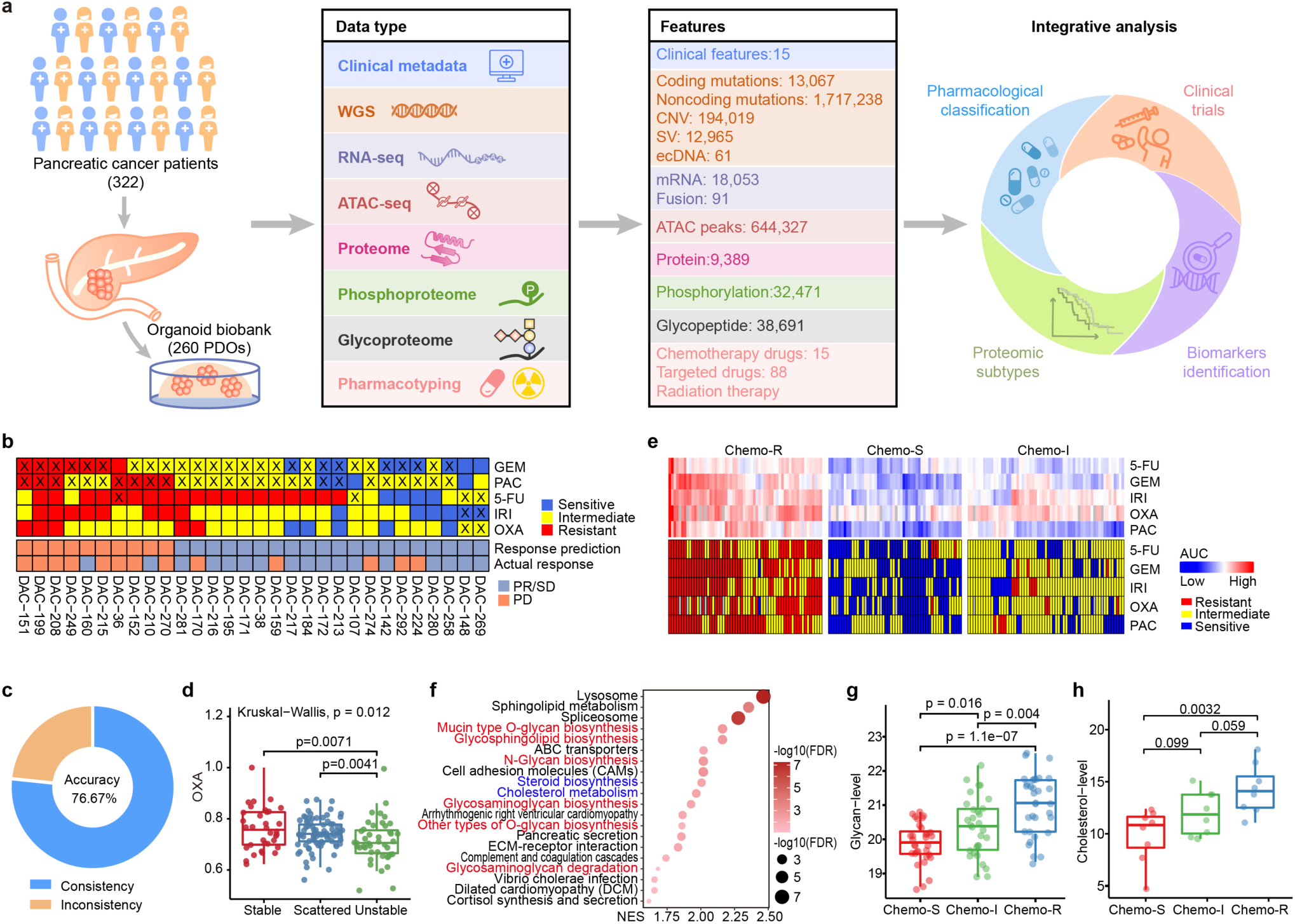
Protein-glycosylation and cholesterol contribute to the chemotherapy resistant. **a**, Schematic diagram showing summary of data and metadata generated in this study. **b**, By comparing the predicted results with the actual treatment responses, the accuracy of the organoid drug sensitivity score in predicting treatment responses was evaluated. Symbol ‘X’ indicated the chemotherapy drugs used in clinical practice of corresponding patients. Using the RECIST v1.1 criteria, patients’ treatment responses are classified as partial responses (PR), stable disease (SD), and progressive disease (PD). Based on the results of organoid drug sensitivity tests, chemotherapy regimen are assigned sensitivity scores (Refer to Table 1). If the total sensitivity score for the chemotherapy regimen is ≥ 2, it predicts a patient response to the regimen; otherwise, it predicts no response. **c**, Overall match rate between organoid drug sensitivity and patient clinical outcomes. **d**, Normalized AUC of OXA among ‘stable’, ‘scattered’ and ‘unstable’ subtypes. P-values are calculated by Kruskal-Wallis test. **e**, Heatmap (up) showing AUC profile of 5 chemotherapeutic drugs which are clustered into three subtypes and heatmap (down) showing subgroup of each chemotherapeutic drug (red: resistant with top 25% AUC; blue: sensitive with lowest 25% AUC; yellow: intermedium). **f**, Top 20 significantly enriched pathways in the Chemo-R subtype. Shown are the normalized enrichment score (NES) and -log10(FDR) of KEGG pathways calculated by GSEA based on ranked proteins. **g**, Comparison of protein glycosylation levels in three pharmacological subtypes at the sample level, differential glycopeptides with p < 0.01 were included in the analysis. P-values are calculated by t-test. **h**, Comparison of total intracellular free cholesterol level (µmol/L per 200,000 cells) in three pharmacological subtypes. P-values are calculated by t-test.

**Table 1.** Organoid drug sensitivity score.

| Single drug sensitivity test results in vitro | Single drug sensitivity score | Sensitivity score for treatment regimen | Prediction of treatment response |
| --- | --- | --- | --- |
| ● Resistant | 0 | Drug A + Drug B +<br>Drug C+ ..... | $\geq 2$ : response |
| ● Median | 1 |  |  |
| ● Sensitive | 2 |  | < 2 : no response |

These 260 organoid lines were divided into a discovery set (197 organoid lines) and a validation set (63 organoid lines). The discovery set underwent comprehensive characterization, including whole-genome sequencing (WGS), RNA sequencing (RNA-seq), assay for transposase-accessible chromatin using sequencing (ATAC-seq), proteomics, phosphoproteomics, glycoproteomics, drug sensitivity screening (Fig. 1a and Supplementary Table 1). The validation set was analyzed using RNA-seq and drug sensitivity testing.

### Increased glycan biosynthesis and cholesterol metabolism pathways in chemoresistant organoids

The efficacy of chemotherapy, a crucial determinant of treatment success, varies significantly among patients^12^. This variability was highlighted through pharmacotyping of 5 standard chemotherapeutic drugs GEM, 5-FU, irinotecan (IRI), oxaliplatin (OXA) and paclitaxel (PAC) used for PDAC patients, which revealed distinct responses among organoids, as assessed via dose-response curves and the corresponding area under the curve (AUC) values (Extended Data Fig. 1a). Notably, pharmacotyping results were consistent across multiple organoid passages (Extended Data Fig. 1b-f).

Organoids were categorized into three subgroups based on their response to each chemotherapeutic agent: resistant (top 25% AUC), sensitive (lowest 25% AUC), and intermediate (middle 50% AUC). To evaluate the clinical relevance of these subgroupings, retrospective clinical follow-up data were obtained for 30 PDAC patients from whose organoids were derived via EUS-FNA. Patients were divided into two groups based on organoid sensitivity and their actual chemotherapy regimens: those predicted to have good response (score ≥ 2) and those predicted to have poor response (score < 2). Clinical outcomes showed 17 cases of partial responses (PR) or stable disease (SD), and 13 cases of progressive disease (PD). Patients in the good response group exhibited favorable clinical outcomes (PR/SD), while those in the poor response group showed adverse outcomes (PD) (Fig. 1b). The overall match rate was 76.67% between organoid sensitivity and patient outcomes (Fig. 1c).

Organoids were further classified into three subtypes based on structural variant distribution: ‘stable’ (lowest 25% structural variations), ‘unstable’ (top 25% structural variations), and ‘scattered’ subtype (intermediate levels) (Extended Data Fig. 1g). Consistent with previous study^13^, the ‘unstable’ subtype demonstrated increased sensitivity to OXA but not to the other four chemotherapy drugs (Fig. 1d, Extended Data Fig. 1h-k).

Clustering analysis of AUC values for the five chemotherapeutic drugs identified three major groups: a chemoresistant group (Chemo-R, resistant to most drugs), a chemosensitive group (Chemo-S, sensitive to most drugs), and an intermediate group (Chemo-I, moderately sensitive to the drugs) (Fig. 1e). Pathway enrichment analysis of proteomic data revealed increased activity in glycan biosynthesis and cholesterol metabolism pathways in the Chemo-R group (Fig. 1f). Glycans, which are covalently bound to proteins or lipids, form glycosylation products such as glycoproteins and glycolipids^14^. Protein glycosylation levels were highest in the Chemo-R group, as determined by the analysis of glycosylation data (Fig. 1g). Similarly, free cholesterol levels were also highest in the Chemo-R group (Fig. 1h). These findings suggest that the elevated protein glycosylation and cholesterol metabolism may play a critical role in driving chemoresistance in pancreatic cancer.

### Proteomic subtyping reveals a high glycosylation subtype

Through proteomics and phosphoproteomics analyses, we quantified 7,225 proteins and 17,781 phosphosites (detected in at least half of the samples). Utilizing K-means clustering on proteins with the highest variability, we identified 3 distinct proteomic subtypes (Extended Data Fig. 2a and Supplementary Table 1). Pathway-centric analysis of these subtypes revealed unique biological characteristics, with one subtype showing significant enrichment in glycan biosynthesis pathways (Extended Data Fig. 2b). Gene set variation analysis (GSVA) further validated this distinct subtype, characterized by high glycan biosynthesis activity across all samples (Fig. 2a). This subtype was designated as the glycan-high cluster (Glycan-H), while the other two subtypes were named Glycan-low (Glycan-L) and Glycan-intermedium (Glycan-I).

**Fig. 2:**
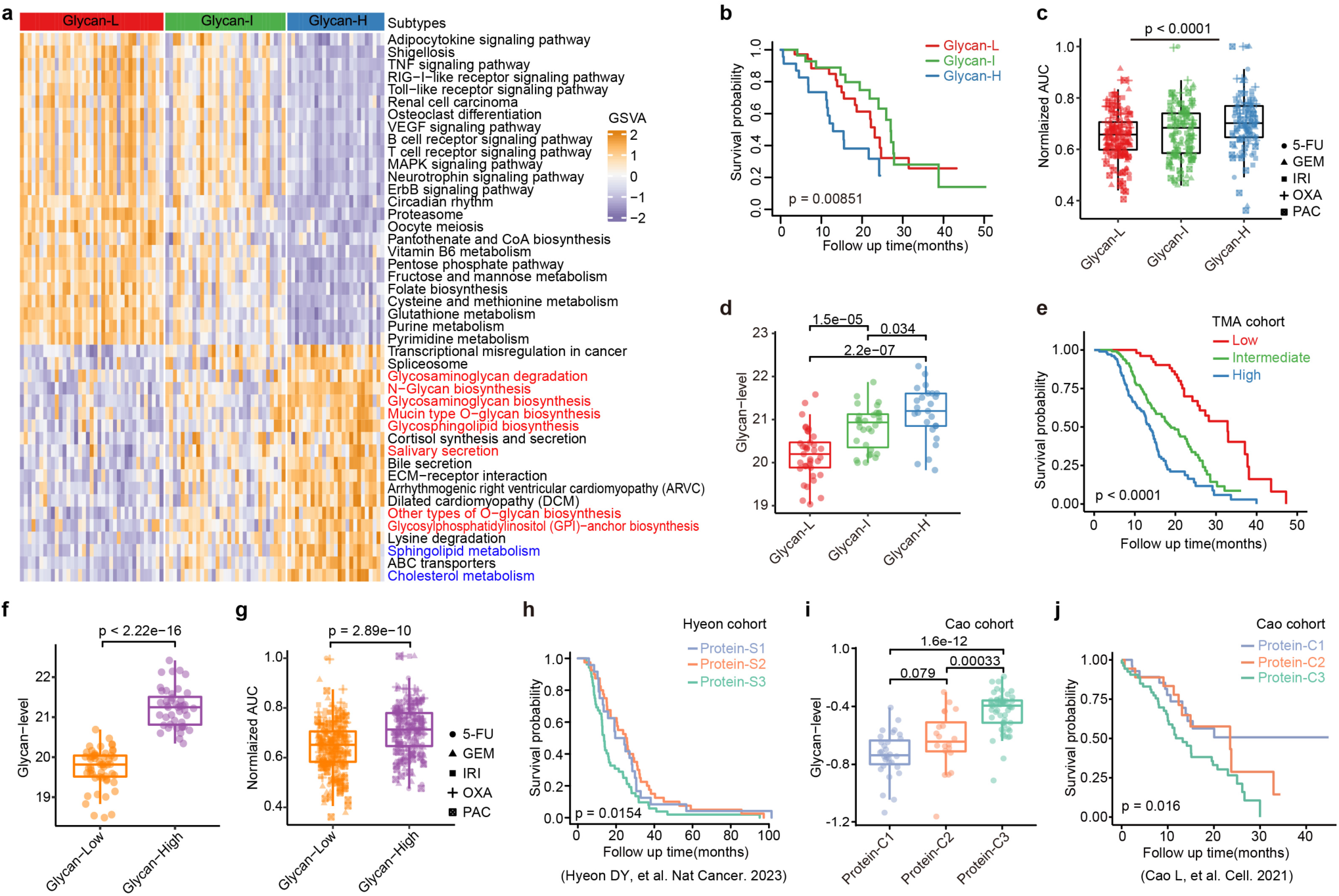
Proteomic subtypes with strong prognostic and chemosensitivity implications. **a**, Heatmap showing pathways with differential GSVA enrichment scores among three proteomic subtypes (ANOVA, FDR<0.01). **b**, Kaplan-Meier survival curves for overall survival (OS) among three proteomic subtypes. Log-rank test is used for statistical analyses. **c**, Normalized AUC of 5 chemotherapeutic drugs among three proteomic subtypes. P-value is calculated by one-sided Wilcoxon rank-sum test (Glycan-H vs Glycan-L). **d**, Comparison of glycan level among three proteomic subtypes at the sample level, differential glycopeptides with p < 0.01 were included in the analysis. P-values are calculated by t-test. **e**, Kaplan-Meier survival curves for OS in Tissue Microarray (TMA) which were divided into three groups based on SNA level. Log-rank test is used for statistical analyses. **f**, Comparison of protein glycosylation levels in the two glycoproteomic subtypes at the sample level, differential glycopeptides with p < 0.01 were included in the analysis. P-value is calculated by t-test. **g**, Normalized AUC values of five chemotherapeutic drugs across the two glycoproteomic subtypes. P-value is calculated by one-sided Wilcoxon rank-sum test. **h**, Kaplan-Meier survival curves for OS among three subgroups of Hyeon cohort. Log-rank test is used for statistical analyses. **i**, Comparison of glycan level among three subgroups of Cao cohort at the sample level, differential glycopeptides with p < 0.01 were included in the analysis. P-values are calculated by t-test. **j**, Kaplan-Meier survival curves for OS among three subgroups of Cao cohort. Log-rank test is used for statistical analyses.

The Glycan-H subtype exhibited the poorest overall survival and the highest resistance to chemotherapy among the three proteomic subtypes (Fig. 2b,c). Furthermore, the cholesterol metabolism pathway was significantly enriched in the Glycan-H subtype (Fig. 2a and Extended Data Fig. 2b), suggesting a potential role for cholesterol metabolism in chemotherapy resistance in PDAC. Notably, basic clinical parameters, such as age, sex, and tumor stage, were similar across the three proteomic subtypes (Extended Data Fig. 2c).

To determine whether the proteomic subtype-specific pathway profiles were reflected at the transcriptomic level, we analyzed the expression of glycan pathway components across the three subtypes. However, we observed discordance between protein and RNA expression levels for most glycan pathway genes in PDAC (Extended Data Fig. 2d,e). The overall correlation between RNA and protein levels was low (r < 0.25) (Extended Data Fig. 2f,g), underscoring the importance of both proteomic and transcriptomic analyses in identifying novel PDAC subtypes.

To validate the proteomic subtypes, we analyzed glycoproteomics data from the organoid biobank. The Glycan-H subtype exhibited a pronounced presence of glycoproteins (Fig. 2d). Differential analysis of glycan categories further confirmed elevated glycoprotein expression in the Glycan-H subtype (Extended Data Fig. 2h). Additionally, we evaluated sialylated glycans in glycoproteins from randomly selected organoids and tissues. Sialylated glycan levels were significantly higher in Glycan-H organoids and tissues compared to the other two subtypes (Extended Data Fig. 2i-l).

To assess the clinical implications of these findings, we conducted tissue microarray (TMA) analyses on patient samples (n = 270) with complete clinical data. Sialylated glycan expression levels were categorized into low, intermediate, and high groups (Extended Data Fig. 2m and Supplementary Table 2). Kaplan-Meier analysis revealed that patients with high sialylated glycans expression had the poorest overall survival (Fig. 2e). Clustering analysis using glycoproteomics data further confirmed that the Glycan-H subtype exhibited significantly higher glycoprotein levels and lower chemosensitivity compared to the Glycan-L subtype (Fig. 2f,g and Extended Data Fig. 2n,o). Consistent with proteomic findings, the Glycan-H subtype was also enriched in cholesterol metabolism pathways (Extended Data Fig. 2o).

To further validate the three proteomic subtypes, we reanalyzed two published PDAC cohorts: the Hyeon cohort^15^ and the Cao cohort^16^. Using the same classification method, we identified three protein subtypes (Protein-S1, Protein-S2, and Protein-S3) in the Hyeon cohort (Extended Data Fig. 3a). Pathway enrichment analysis confirmed that Protein-S3 corresponded to the glycan-high subtype. Consistently, Protein-S3 exhibited the poorest prognosis in the Hyeon cohort (Fig. 2h). Similarly, the Cao cohort was stratified into three subtypes based on this signature (Extended Data Fig. 3c,d). Patients in the Protein-C3 subtype, which exhibited the highest levels of protein glycosylation, had the worst overall survival compared to the other two subtypes (Fig. 2i,j). Notably, the glycan-high subtypes in both cohorts were also enriched in cholesterol metabolism pathways (Extended Data Fig. 3a,b,d,e). These findings highlight the clinical and biological significance of the glycan-high subtype, which is characterized by elevated protein glycosylation, enrichment in cholesterol metabolism pathways, poor prognosis, and chemotherapy resistance in PDAC.

### Pharmacogenomic insights of PDAC

Although translating genomic data into actionable therapeutic strategies holds promise for cancer treatment^12^, it remains a significant challenge in the clinical management of pancreatic cancer. To evaluate the clinical relevance of genomic variants identified in our cohort, we applied OncoKB and CGI evidence tiers to systematically assess clinically actionable genomic variants (AGVs) across all tumors (pan-cancer) (Extended Data Fig. 4a and Supplementary Table 3). Our analysis revealed that 79.6% of samples harbored AGVs of clinical significance, potentially influencing treatment decisions involving targeted therapies (Extended Data Fig. 4b).

To explore therapeutic opportunities, we conducted drug sensitivity testing in organoids targeting key pathways implicated in pancreatic cancer, including RTK, MAPK, PI3K-AKT-mTOR signaling, DNA damage repair, cell cycle regulation, and epigenetic modifications. Broad-spectrum multi-kinase inhibitors were included due to their efficacy against treatment-resistant and aggressive PDAC subtypes^17^. Chemotherapeutic drug selection adhered to NCCN guidelines. Given the increased glycan biosynthesis and cholesterol metabolism activity observed in Chemo-R organoid lines (Fig. 1f), we also tested the sensitivity of glycosylation inhibitors (Benzyl-α-GalNAc, Lith-O-ASP, SGN-2FF, and Tunicamycin) and cholesterol metabolism inhibitors (Avasimibe, Ezetimibe, Nevanimibe, NB-598, Atorvastatin, Lovastatin, Pitavastatin, and Simvastatin) within our cohort.

A comprehensive therapeutic profile was established for 184 PDAC organoids, evaluating their responses to 88 targeted agents and 15 chemotherapeutic drugs (Extended Data Fig. 4c and Supplementary Table 3). Among these, 14 targeted drugs, 8 mono-chemotherapeutic agents, and 8 combined chemotherapeutic agents currently undergoing clinical trials or having completed clinical trials for PDAC. Rigorous quality control ensured high correlation among biological replicates across separate screenings (Extended Data Fig. 4d). Drug efficacy was quantified using IC50 (the concentration reducing viability by 50%) and AUC values (Supplementary Table 3), with both metrics showing strong correlation and consistency (Extended Data Fig. 4e). AUC values were used uniformly for all analyses.

To assess the translational relevance of organoid therapeutic profiling, we analyzed the alignment between drug sensitivity and corresponding AGVs. Positive outcomes were observed when the sensitivity of organoids with specific alterations fell below the cohort average (Extended Data Fig. 4f), validating 37.68% of AGVs-drug correlations through organoids therapeutic profiling (Extended Data Fig. 4g). When drugs were categorized by their primary targets into six signaling pathways, the strongest correlation between AGVs and drug sensitivity was observed for compounds targeting the PI3K, MAPK, and DNA damage repair pathways, while the weakest correlation was noted for RTK and epigenetic pathway inhibitors (Extended Data Fig. 4h). This suggests that genomic alterations in PI3K and MAPK pathways may confer resistance to upstream RTK inhibitors. This hypothesis is supported by previous findings showing that EGFR-targeting compounds are effective in cells with *EGFR* and *ERBB2* alterations but not in those with downstream *ERK-MAPK* mutations, such as mutant *RAS*^18^.

Interestingly, we observed discrepancies in sensitivity to ERBB2 inhibitors among organoid lines with and without *ERBB2* amplifications (Extended Data Fig. 4i). Elevated *IGFBP4* expression was strongly associated with sensitivity to the ERBB2 inhibitor lapatinib (Extended Data Fig. 4j,k), aligning with previous studies highlighted IGFBP4’s inhibition of RTK and AKT signaling pathway^19^. This led us to hypothesize that reduced sensitivity o to lapatinib in *ERBB2*-amplified organoids might be linked to a concurrent dependency on IGFBP4 (Extended Data Fig. 4l).

### Statins overcome intrinsic chemoresistance in PDAC

To discover therapeutic agents that could effectively target chemoresistant PDAC, we conducted a sensitivity comparison between Chemo-R and Chemo-S organoid lines using the drug library described above. This analysis revealed that Chemo-R organoids were resistant to most standard single-agent and combination chemotherapy regimens currently used in clinical practice, such as AG and FFX (Fig. 3a). Surprisingly, HMGCR inhibitors (Atorvastatin, Lovastatin, Pitavastatin, and Simvastatin) and the acyl coenzyme A-cholesterol acyltransferase inhibitor Avasimibe demonstrated significantly higher efficacy in Chemo-R organoids compared to Chemo-S organoids (Fig. 3a).

**Fig. 3:**
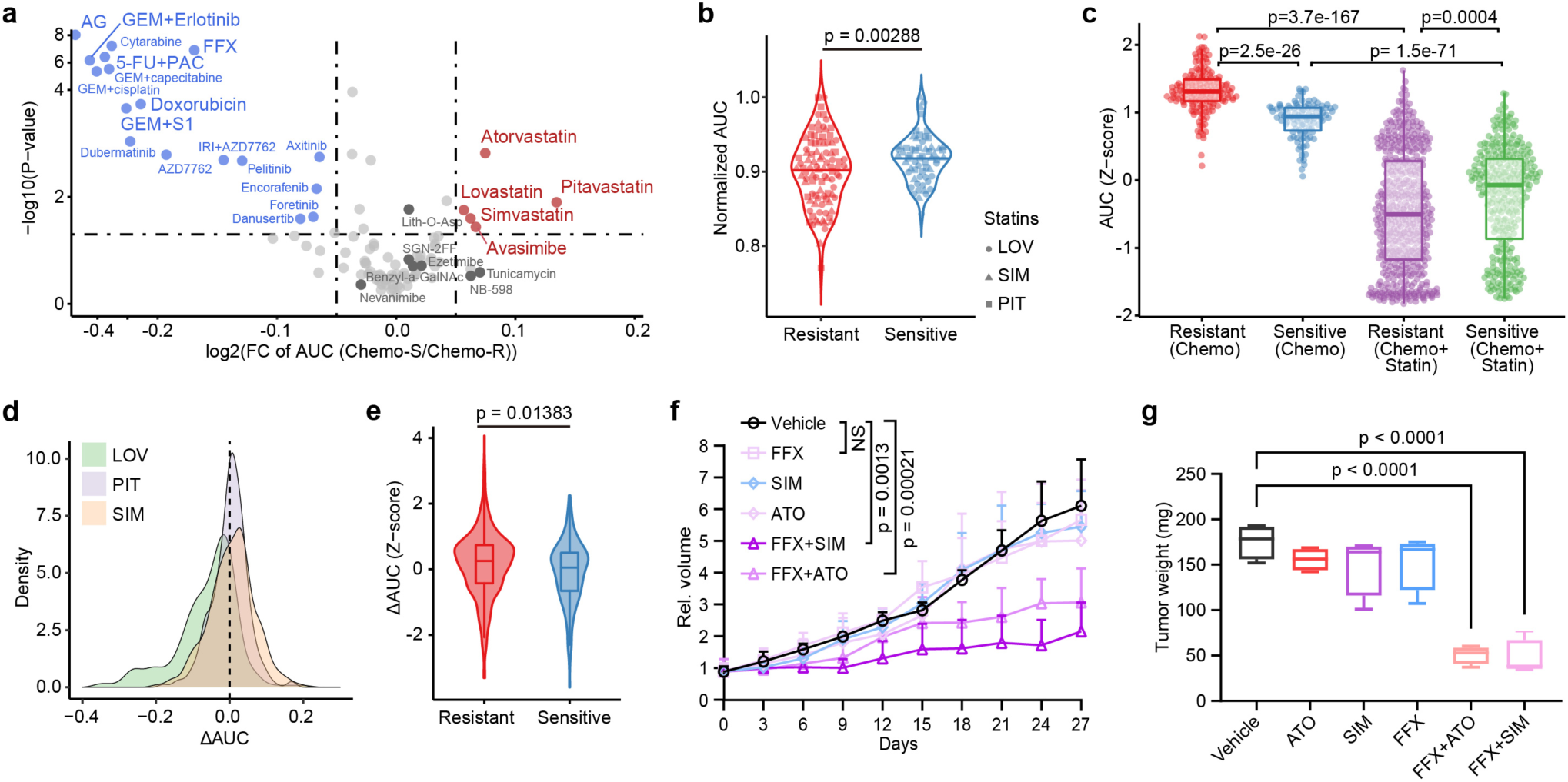
Statins overcome intrinsic chemoresistance in PDAC. **a**, Volcano plot showing the distribution of inhibitors in Chemo-R and Chemo-S organoids. Statins and Avasimibe with preferential inhibition in Chemo-R organoids are shown in red. Four protein glycosylation inhibitors and other three cholesterol metabolism inhibitors are labeled in dark gray. **b**, Normalized AUC of 3 statins among the two chemotherapeutic subtypes in validation set. P-value is calculated by one-sided Wilcoxon rank-sum test. The abbreviation of the inhibitors are as follows: Lovastatin (LOV), Pitavastatin (PIT), and Simvastatin (SIM). **c**, AUC (Z-score) comparison for chemotherapy only and combination of chemotherapy and statins among the two chemotherapeutic subtypes in validation set. P-value is calculated by one-sided Wilcoxon rank-sum test. **d**, Density distribution of ΔAUC across all combination treatment in validation set. **e**, Scaled ΔAUC distribution among the two chemotherapeutic subtypes in validation set. P-value is calculated by one-sided Wilcoxon rank-sum test. **f**, Tumor volume of DAC-71 derived subcutaneous xenografts (n = 6 per group) in SCID mice following treatment with the indicated drugs. P-values are calculated by t-test. Atorvastatin is abbreviated as ‘ATO’. **g**, Boxplot showing tumor weight changes of DAC-71 derived orthotopic xenografts for each group with indicated drug treatment. P-values are calculated by t-test.

Since stains exhibited the most significantly suppression activity in Chemo-R organoids, we further confirmed the therapeutic potential of statins in chemoresistant PDAC organoids. Cell viability assays were performed using chemotherapy alone or in combination with statins in the validation set of 52 organoid lines. These organoids were categorized into Sensitive and Resistant groups based on their response to five standard chemotherapy drugs (Extended Data Fig. 5a). The Resistant group displayed greater sensitivity to statins compared to the Sensitive group (Fig. 3b). Moreover, combining chemotherapy with statins significantly enhanced the inhibition of cell viability, particularly in the Resistant group (Fig. 3c and Extended Data Fig. 5b).

To evaluate the synergy between statins and chemotherapy, we compared the observed response of organoids to combination therapy with the expected response calculated using the Bliss independence model, which predicts combination effects based on monotherapy activities. Combination therapy pairs (3 statins × 5 chemotherapy drugs = 15 pairs) were tested across all 52 organoid lines, resulting in 780 organoid-combination pairs. Synergy was defined as a reduction in the actual combination AUC greater than predicted by the Bliss model (ΔAUC > 0). Remarkably, 43.59% of the organoid-combination pairs exhibited synergy (Fig. 3d), with a higher likelihood of synergistic effects observed in Resistant organoids (Fig. 3e and Extended Data Fig. 5c).

To assess the chemosensitizing effects of statins *in vivo*, we used a Chemoresistant PDAC organoid line (DAC-71) for orthotopic and subcutaneous transplantation into immunodeficient SCID mice. Combined treatment with statins and FFX significantly inhibited tumor growth compared to chemotherapy alone in both orthotopic and subcutaneous transplantation models (Fig. 3f,g and Extended Data Fig. 5d).

### Chemoresistant PDAC is associated with increased EMT signatures

Emerging evidence suggests that metabolic reprogramming plays a critical role in chemotherapy resistance^20^. We further analyzed the correlation between metabolic pathways and the chemotherapy sensitivity or statin treatment using our proteomic dataset. Interestingly, the metabolic pathways correlated with chemotherapy resistance and statin sensitivity were inversely related. Specifically, glycan and lipid metabolism pathways, including cholesterol metabolism, were positively associated with chemotherapy resistance but also with increased sensitivity to statins (Fig. 4a).

**Fig. 4:**
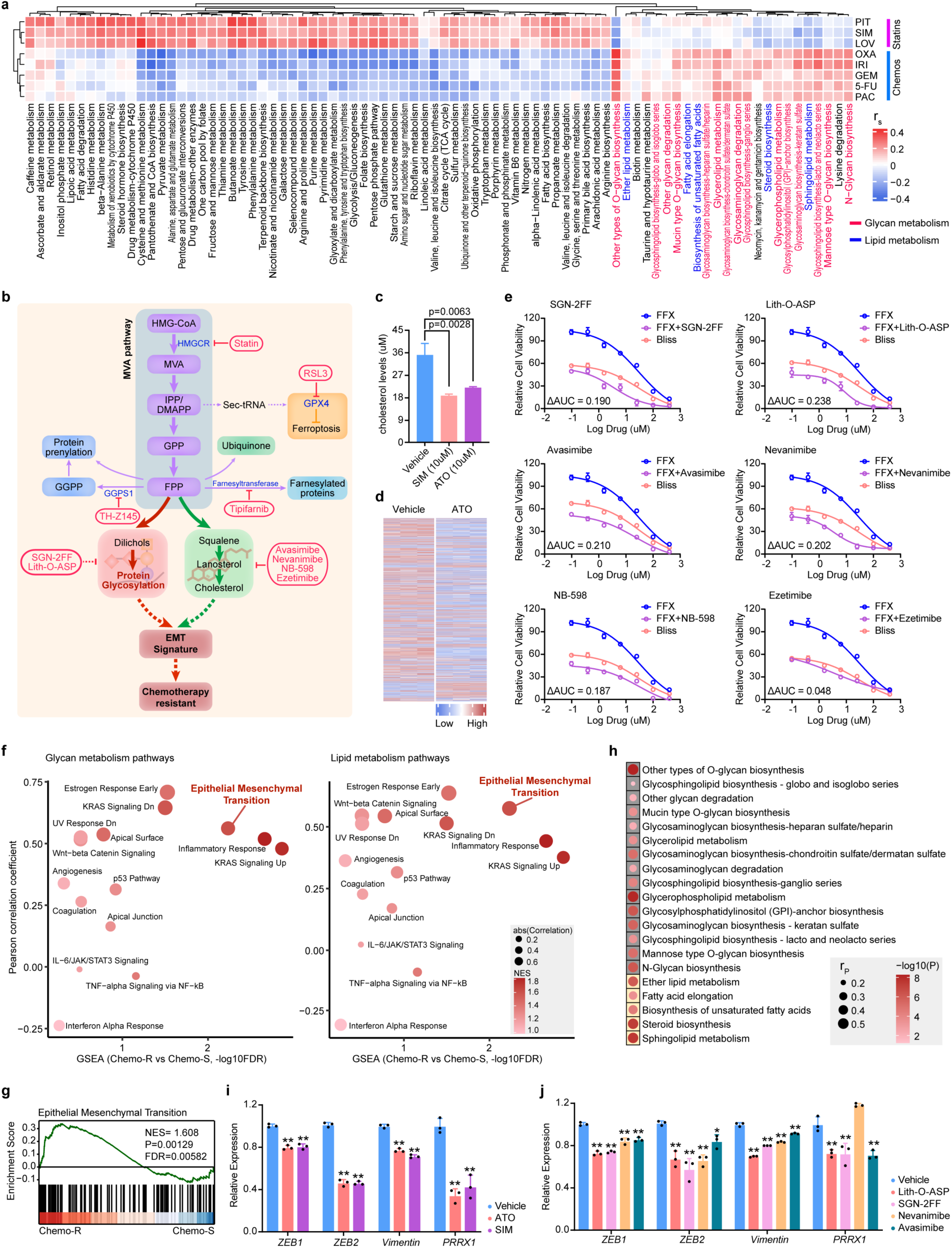
Mechanism of statin-induced increased chemotherapy sensitivity in PDAC. **a**, A heatmap displaying the Spearman correlation between GSVA enrichment scores of metabolic pathways derived from proteomic profiles and the AUC values of the indicated drugs. Metabolic pathways were obtained from the KEGG database. **b**, Schematic representation of the mevalonate (MVA) and downstream pathways, highlighting specific inhibitors targeting the pathways downstream of the MVA which have been tested. **c**, Change of intracellular free cholesterol level (µmol/L, per 1,000,000 cells) in DAC-36 organoid lines following statin treatment. P-values are calculated by t-test. **d**, Heatmap showing glycosylation sites significantly changed between ATO treated and control organoid line (DAC-38). **e**, Dose response curves of FFX in combination with glycosylation and cholesterol metabolism inhibitors in the DAC-36 organoid line. The IC50 concentrations of glycosylation and cholesterol metabolism inhibitors were used to treat organoids. **f**, Bubble chart showing hallmark pathways significantly enriched in Chemo-R and their correlation with glycan metabolism (left) or lipid metabolism (right). The x-axis represents GSEA enrichment significance for gene sets in Hallmark of Chemo-R versus Chemo-S. The y-axis represents Pearson correlation coefficients between GSVA enrichment scores of gene sets in Hallmark and genes in protein glycosylation or lipid metabolism related pathways derived from proteomic data. **g**, GSEA enrichment plot for EMT pathway in Chemo-R versus Chemo-S. **h**, Correlation plot between EMT pathway and glycan or lipid metabolism related pathways. Correlation coefficients (r_P_) are calculated by Pearson correlation analysis based on GSVA enrichment scores of EMT pathway and genes in glycan and lipid metabolism pathways derived from protein data. **i-j**, Bar plot showing the changes in expression levels for EMT related genes after treatment with statins (**i**), protein glycosylation inhibitors and cholesterol metabolism inhibitors (**j**) in the DAC-36 organoid line. Simvastatin (SIM), Atorvastatin (ATO). The IC20 concentrations of inhibitors were used to treat organoids.

Statins are known to reduce cholesterol levels by inhibiting HMGCR in the mevalonate (MVA) pathway which is an essential metabolic pathway for tumor growth^21^ (Fig. 4b). Enrichment analysis of MVA downstream pathways revealed significant enrichment of glycan and cholesterol metabolism in Chemo-R PDAC organoids compared to Chemo-S organoids (Extended Data Fig. 6a,b). Whereas, other MVA downstream processes, such as farnesyltransferase-mediated protein farnesylation, ferroptosis, protein geranylgeranylation, and ubiquinone metabolism, were not significantly enriched in Chemo-R organoids (Extended Data Fig. 6c-f).

As expected, statins treatment resulted in a significant decrease in cellular cholesterol content (Fig. 4c). Meanwhile, treatment with ATO led to a marked reduction in protein glycosylation levels in chemoresistant organoids (Fig. 4d). Differential analysis across four glycan categories revealed a significant suppression of glycoprotein levels, with the most pronounced decrease observed in sialic acid levels (Extended Data Fig. 6g). This was further confirmed by reduced sialic acid immunofluorescence staining in atorvastatin-treated organoids (Extended Data Fig. 6h). Similarly, statin administration significantly reduced protein glycosylation levels in chemoresistant DAC-71 xenografts (Extended Data Fig. 6i). These findings suggest that statins suppress the MVA pathway and enhance chemotherapy sensitivity by inhibiting glycan and cholesterol metabolism pathways.

To further evaluate chemotherapy sensitivity in chemoresistant organoids, we inhibited downstream signaling of the MVA pathway using specific inhibitors (Fig. 4b). Inhibitors targeting protein glycosylation (SGN-2FF and Lith-O-ASP) and cholesterol metabolism (Avasimibe, Nevanimibe, NB-598, and Ezetimibe) exhibited synergistic effects when combined with FFX treatment (Fig. 4b,e). In contrast, inhibitors targeting other MVA downstream pathways, such as farnesyltransferase (Tipifarnib), ferroptosis (RSL3), and protein geranylgeranylation (TH-Z145), did not show synergistic effects (Fig. 4b and Extended Data Fig. 6j).

To investigate the downstream mechanisms of glycan and cholesterol metabolism in regulating chemotherapy resistance, we performed gene set enrichment analysis (GSEA) and analyzed their correlation with protein glycosylation and lipid metabolism pathways in Chemo-R versus Chemo-S PDAC organoids. The epithelial-to-mesenchymal transition (EMT) signaling pathway was significantly enriched in Chemo-R organoids and strongly correlated with glycan and cholesterol metabolism pathways (Fig. 4f,g). Additionally, various glycan and lipid metabolism pathways were significantly associated with EMT (Fig. 4h).

EMT has been identified as a fundamental mechanism of chemotherapy resistance^22^. To confirm the link between glycan and cholesterol metabolism and EMT signatures, we treated chemoresistant PDAC organoids with upstream HMGCR inhibitors (atorvastatin and simvastatin) as well as downstream inhibitors of protein glycosylation (Lith-O-ASP and SGN-2FF) and cholesterol metabolism (Avasimibe and Nevanimibe). All tested inhibitors significantly suppressed the expression of EMT signature genes, including *ZEB1*, *ZEB2*, *Vimentin*, and *PRRX1* (Fig. 4i,j). In contrast, inhibitors targeting other MVA downstream pathways (Tipifarnib, RSL3, and TH-Z145) did not reduce EMT signature gene expression (Extended Data Fig. 6k). These results suggest that statins effectively inhibit EMT in chemoresistant PDAC organoids by attenuating glycan and cholesterol metabolism, providing a mechanistic basis for their chemosensitizing effects.

### Statins improve chemotherapy efficacy in clinical treatment

To investigate the efficacy of statins in improving outcomes for patients with advanced chemoresistant pancreatic cancer, we conducted a single-arm, phase 2 clinical trial (NCT06241352). This study enrolled 34 patients with locally advanced or metastatic pancreatic cancer, all of whom had reached a plateau phase in their tumor markers (CA19-9 or CEA, with CA19-9-negative patients referenced against CEA) following prior chemotherapy. During the treatment, the stabilization of tumor markers at a plateau is often considered an indication that the tumor is developing resistance to therapy. By adding statins to the existing treatment regimen, we aim to validate whether statins can alleviate patients’ resistance to chemotherapy. Patients received their standard chemotherapy regimen alongside atorvastatin 80mg/day. Atorvastatin treatment was discontinued if tumor marker levels increased by more than 20% above baseline (Fig. 5a).

**Fig. 5:**
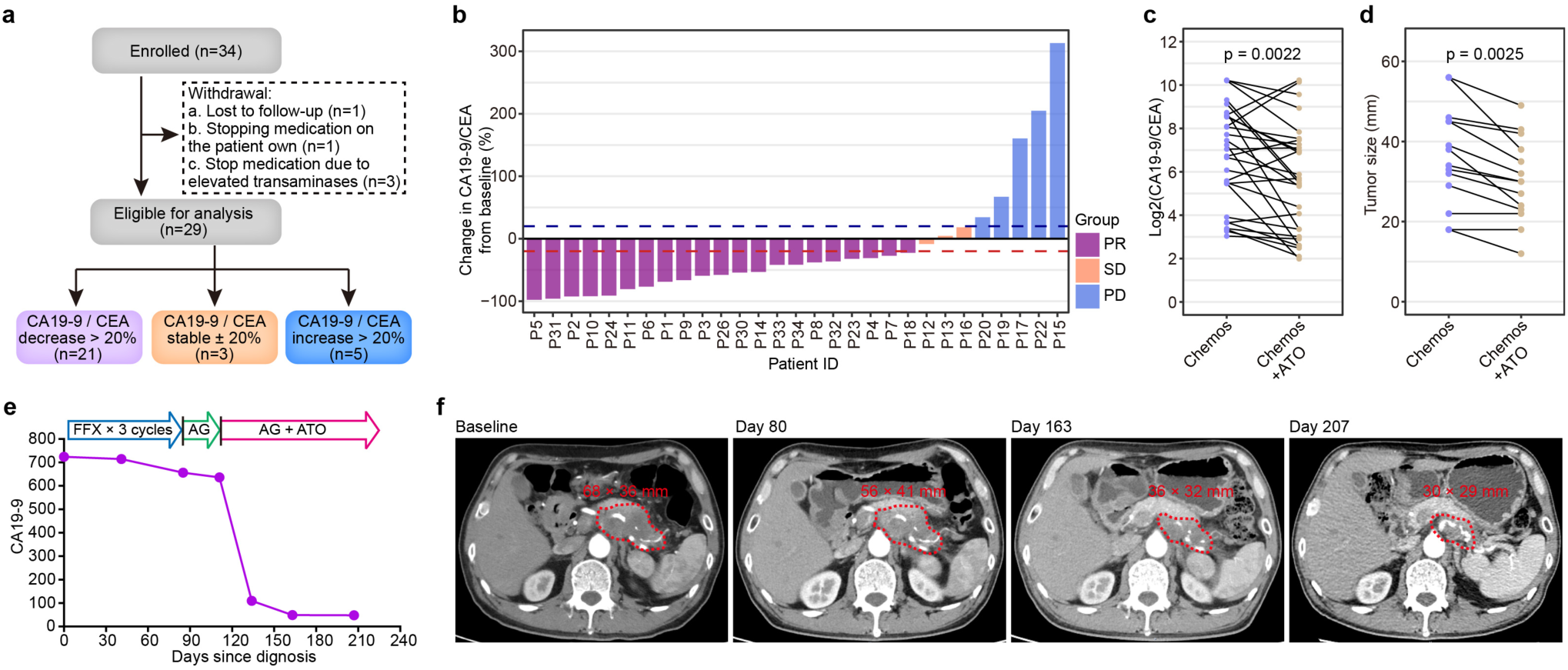
Statins improve chemotherapy efficacy in clinical treatment. **a**, Schematic of clinical trials including patient enrollment and group allocation based on treatment efficacy. **b**, Bar plot showing the changes of CA19-9 or CEA in 29 patients after taking atorvastatin for at least one month. The CA19-9 level of the patient before taking Atorvastatin (ATO) was used as the baseline. **c-d**, Line plot showing a significant decrease in CA19-9 or CEA levels (**c**) and tumor size (**d**) after taking ATO for one month. P-values are calculated by Wilcoxon rank-sum test. **e**, Serum CA19-9 measurements from diagnosis throughout the patient’s (P2) treatment course. The arrow indicates transition from FFX chemotherapy to AG chemotherapy to AG and statin combine therapy. **f**, Representative radiation examination images for the patient’s (P2) at different time points. The red circles mark the tumor positions.

Of the 34 patients enrolled, one was lost to follow-up, one voluntarily discontinued medication, and three discontinued treatments due to elevated transaminases. Among the remaining 29 patients, preliminary results showed that 21 patients (72.4%) experienced a greater than 20% decrease in tumor markers (CA19-9 or CEA) one month after the addition of atorvastatin (Fig. 5b and Extended Data Fig. 7a, Supplementary Table 4). Based on the follow-up results obtained so far, the 21 patients experienced an average of 66.86 days with tumor marker levels below baseline. Notably, two of these patients (Patient #3 and Patient #18) initially responded to the combination therapy but later developed resistance, with tumor markers rapidly increasing again. Three patients (Patient #12, Patient #13 and Patient #16) exhibited stable tumor marker levels, while five patients (Patient #15, Patient #17, Patient #19, Patient #20 and Patient #22) showed an increase in tumor markers despite the combination therapy (Fig. 5b and Extended Data Fig. 7b).

Overall, patients with advanced pancreatic cancer in the plateau phase of chemotherapy demonstrated significant reductions in tumor markers and tumor size following the addition of atorvastatin to their chemotherapy regimen (Fig. 5c,d). For instance, Patient #2 had reached the plateau stage after three cycles of FFX and AG treatment, as indicated by CA19-9 levels fluctuating by less than 20% over a period of more than four months. The combination of atorvastatin and AG treatment resulted in impressive responses, with significant reductions in both tumor markers and tumor size (Fig. 5e,f).

These results not only exemplify the promising therapeutic role of atorvastatin in enhancing chemotherapy response but also suggest a potential paradigm shift in the management of pancreatic cancer, warranting further investigation into the use of statins combined with standard chemotherapy in this challenging disease.

## Discussion

In this study, we conducted a comprehensive proteogenomic analysis of PDAC by integrating multi-omic data to uncover therapeutic vulnerabilities and classification strategies. By correlating multi-omics profiles with drug sensitivity data, we identified that chemoresistant PDAC subsets shared a distinct signature characterized by elevated glycosylation and cholesterol metabolism activity. Furthermore, we demonstrated that statins significantly inhibited the growth of chemoresistant PDAC organoids. Notably, in a phase 2 clinical trial (NCT06241352), combining atorvastatin with chemotherapy for advanced pancreatic cancer resulted in remarkable reductions in both tumor markers and tumor size.

EMT and metabolic reprogramming are increasingly recognized as fundamental mechanisms underlying chemotherapy resistance in cancer, including PDAC^23^. Chemotherapy-resistant PDAC cells often exhibit elevated expression of EMT markers, including vimentin and N-cadherin, alongside reduced expression of epithelial markers such as E-cadherin^24^. Cholesterol metabolism, mediated by the MVA pathway, is essential for membrane synthesis, signaling, and energy production in cancer cells. Elevated cholesterol metabolism has been shown to support EMT by modulating membrane fluidity and lipid raft formation, which are critical for the activation of EMT-related signaling pathways^21^. Additionally, increased glycosylation of surface proteins and signaling molecules in chemoresistant PDAC has been linked to the activation of EMT and other pro-survival pathways^14^. These studies underscore the importance of developing combination therapies that target both EMT and metabolic pathways to improve outcomes for patients with chemoresistant cancers.

By generating a patient-derived pancreatic cancer organoid biobank and integrating multi-omics profiling, we identified elevated glycosylation and cholesterol metabolism as key features associated with enhanced EMT signatures in chemoresistant PDAC. Our results showed that statins were among the most effective agents in inhibiting the growth of chemoresistant pancreatic cancer organoids. Consistently, combining statins with chemotherapy demonstrated significant therapeutic efficacy in organoid models both *in vitro* and *in vivo*. Mechanistically, our findings suggest that statins enhance chemotherapy sensitivity by reducing protein glycosylation and inhibiting EMT signatures in organoids. Abnormal glycosylation has been shown to promote EMT, facilitating tumor progression and drug resistance, ultimately contributing to tumor aggressiveness and therapy failure^22,25,26^. Excitingly, the clinical trial (NCT06241352) combining atorvastatin with chemotherapy in patients with advanced chemoresistant pancreatic cancer demonstrated notable therapeutic responses, with significant reductions in tumor markers and tumor size.

This study provides a detailed proteogenomic and drug sensitivity profile of PDAC, leveraging a substantial cohort of organoid models to highlight the limitations of current reliance on mono- or combination chemotherapy as the standard treatment for PDAC. Importantly, this work offers a valuable resource for understanding PDAC tumor biology and represent a significant step toward developing more effective, personalized treatment strategies. By addressing the challenges of chemoresistance, our findings demonstrate that combining statins with chemotherapy represents a promising and actionable therapeutic approach with significant clinical implications for chemoresistant pancreatic cancer.

## Acknowledgments

We thank Yaqin Yan, Shujue Lan, Yunqing Ci and Ming Chen for providing technical help at the SIBCB Core Facility. We thank Dr. Baozhen Shan, Dr. Wenting Li, Dr. Yuxian Xu and Dr. Yan Xu from Bioinformatics Solutions Inc. for the help of glycoproteomic data analysis. This study was supported by the National Key Research and Development Program of China (No.2020YFA0509000, 2023YFC2506400), the National Science Fund for Distinguished Young Scholars (32125013), the National Natural Science Foundation of China (82303257, 82303575, 82172712, 82172589, 81972913, 81672830, 92253304, 12025107), the Basic Frontier Science Research Program of Chinese Academy of Sciences (ZDBS-LY-SM015), the Shanghai Science and Technology Committee (21XD1424200 and 21ZR1470100), the Shanghai Municipal Science and Technology Major Project, the Innovative Research Team of High-level Local Universities in Shanghai (SHSMUZDCX20211800), and Shanghai Science and Technology Innovation Action Plan (23Y41900200). We acknowledge funding from the National Key Research and Development Program of China (2022YFA1004800), and CAS Project for Young Scientists in Basic Research (YSBR-077).

## Author Contributions

D.G. conceived and designed the experimental approach. Y.G.L., H.W. and Y.H.Z. performed most experiments. S.J.T., Y.R.L., L.N.C., Y.W. and Y.X.M contributed to the computational analysis and statistical analysis. G.J. and S.W.G provided clinical data. H.Z., H.W.Z and X.J.L. provided proteomic, phosphoproteomic and glycoproteomic technical support and data analysis. J.H., Y.K.L., Y.Z., X.H.S., C.L.Z., Y.Q.Z., Y.J., and M.Y.S. helped with the experiments and provided technical support. Y.G.L., S.J.T and D.G. prepared the manuscript as senior authors.

## Competing Interests Statement

The authors have no competing interests to declare.

## Methods

### Patients’ cohort

PDAC tissue samples were collected at Changhai Hospital. All patients involved in this study gave informed consent for the use of their clinical data and surgical specimens and agreed to the release of clinical information that could identify individuals. The protocols for this study were in conformity with national guidelines and received approval from Changhai Hospital’s ethics committee (approval no. CHEC2018-111). Detailed clinical information of these patients was listed in Supplementary Table 1.

### Organoid culture

Tissues from PDAC patients were washed in basic medium (1640 basic medium, 100 µg/ml Primocin, 10 µM Y-27632) for twice and minced into 1 mm^3^-fragments. The tissue fragments were digested with collagenase II (5 mg/ml, Gibco) on an rotator at 37°C for a maximum of 1 hours. The digested tissue supernatant was strained over a 70 µm filter before centrifugation at 1700 rpm. Isolated cells were embedded in droplets of Matrigel Growth Factor Reduced (Corning) and human PDAC medium for PDAC organoids. Organoid culture media: basic medium (advanced DMEM/F12, 10 mM HEPES, 1X GlutaMAX-I, 100 µg/ml Primocin, 1X penicillin/streptomycin solution) and complete medium (advanced DMEM/F12, 10 mM HEPES, 1X GlutaMAX-I, 100 µg/ml Primocin, 1X penicillin/streptomycin solution, 30% Wnt3A conditioned medium, 2% R-spondin conditioned medium, 4% Noggin conditioned medium, 500 nM A83-01, 10 µM Y-27632, 1.56 mM N-acetylcysteine, 10 mM nicotinamide, 10 ng/ml FGF10, 1X B27 supplement, 10 µM forskolin,). The media used for organoid cryopreservation were composed of the corresponding culture medium (90%) and 10% DMSO. The established organoids were routinely tested for mycoplasma contamination. All organoid experiments were performed at the Shanghai Institute of Biochemistry and Cell Biology.

### Animals

Female SCID mice of 6-week-old were purchased from Biocytogen Pharmaceuticals (Beijing) Co., Ltd. All animal work was conducted in accordance with a protocol approved by the Institutional Animal Care and Use Committee (IACUC) of the Center for Excellence in Molecular Cell Science (CEMCS), and ethical approval was received from the IACUC of CEMCS. Mice were bred in specific pathogen free (SPF) animal house with 28°C and 50% humidity. Indicated organoids were inoculated into subcutaneous of the eight-week-old SCID mouse (n = 3 per group, 2X10^6^ cells/injection). After the xenografts became palpable (∼200mm^3^), mice were randomized to experimental groups and control groups, and treated with vehicle, FFX (5-FU 12.5mg/kg, Leucovorin 2.09mg/kg, IRI 0.78mg/kg, OXA 0.45mg/kg), SIM (25 mg/kg), ATO (25 mg/kg), combined administration of FFX and SIM, combined administration of FFX and ATO. Tumor size was measured every 3 days and the tumor volume was calculated with the equation *V* (in mm^3^) = 0.5 X length X width.

### Somatic mutation calling and annotation

Whole genome sequencing (WGS) data were treated according to the Genome Analysis Toolkit (GATK, https://software.broadinstitute.org/gatk/) best practices workflow^27^. First, raw fastq data were processed with Trimmomatic (v0.39)^28^ for adapter trimming and low quality reads filtering and then were aligned to hg19 human genome reference using BWA-mem (v0.7.15)^29^. Samtools (v1.4)^30^ was used to convert the resulting SAM files to compressed BAM files and then sort the BAM files. PCR duplicates were marked with Picard (https://broadinstitute.github.io/picard), and base quality scores were recalibrated using BaseRecalibrator tool of GATK (version 4.0.9.0). Mutect2 was run to call somatic mutations from the tumor-normal paired bam files^27^. In addition, each normal file was conducted with tumor-only mode of Mutect2, then creating a panel of normal (PON) file to filter out expected artifacts and germline variations. The resulted VCF files were annotated with ANNOVAR software and variations with allele frequency less than 0.05 were filtered out^31^. Cancer-related genes were identified by COSMIC and OncoKB database^32,33^.

OncoKB (v3.16) and Cancer Genome Interpreter (CGI)^34^ database were used to identify AGVs in our cohort. The levels of potential therapies range from FDA-approved drugs (level 1), standard care (level 2), clinical evidence (level 3) and biological evidence (level 4).

### Copy number analysis

Tumor-normal paired bam files (sorted, indexed and duplicate marked) were processed with CNVKit^35^ to call somatic copy number variation (CNV) and then GISTIC2.0^36^ (Genomic Identification of Significant Targets in Cancer v2.0.23) was used to identify focal gain and loss regions. CNV amplification (gain) and CNV deletion (loss) per gene referred to ‘2’ and ‘-2’ value in “all_thresholded.by_genes.txt” file generated from GISTIC2.0.

### Tumor mutational burden (TMB) and chromosomal instability (CIN)

Nonsynonymous somatic mutations (including single nucleotide substitutions and indels) were counted and then divided by the size of the coding region (∼35M) to calculate tumor mutation burden (TMB). For calculating the chromosome instability (CIN), we used a weighted-sum approach following another study^37^. Absolute log_2_ ratios of all CNV segments within a chromosome were weighted by the segment length and summed up to derive the instability score for the chromosome. The genome-wide chromosome instability index was derived by summing up the instability score of all 22 autosomes. Wilcoxon rank-sum test was used to calculate difference of TMB and CIN between different conditions.

### Structure variation detection

Somatic structure variations (SVs) including tandem duplications, deletions, inversions and translocations were detected using Delly (v0.8.5)^38^. Sorted, indexed and duplicate marked bam files (tumor-normal paired) were used for input of Delly and the output SVs were annotated with AnnotSV program^39^.

### Germline variant calling

Germline variant calling was performed with HaplotypeCaller of GATK in single-sample mode on both tumor and normal bam files (base quality scores recalibrated). The resulted VCF files were annotated with ANNOVAR software^31^. All resulting variants were limited to the coding region and only variants detected in both tumor and normal samples were remained. Additionally, we filtered out all variants with ≥ 0.01 frequency in ExAC^40^ (East Asian) and the 1000 Genomes Project^41^ (East Asian). To further filter cancer associated germline mutations, we selected variants annotated to pathogenic/likely pathogenic in CLNSIG or InterVar database, and annotated to damaging in at least two deleteriousness prediction methods among SIFT^42^, Polyphen2^43^, LRT^44^, MutationTaster^45^, MutationAssessor^46^ and FATHMM^47^.

### Extrachromosomal circular DNA detection

AmpliconArchitect was used to detect ecDNA based on CNVKit output “sample.cns” files with default parameters as described in the documentation (https://github.com/virajbdeshpande/AmpliconArchitect).^48^ Then AmpliconClassifier was used to classifies the outputs of AmpliconArchitect (https://github.com/jluebeck/AmpliconClassifier)^49^.

### RNA-seq data processing

Raw RNA-seq data were processed with Trimmomatic (v0.39)^28^ for adapter trimming and low quality reads filtering and then were aligned to hg38 human genome reference using STAR (v2.6.0)^50^. Genes expressed as zero in more than 70% samples were filtered out. FPKM (Fragments Per Kilobase of transcript per Million mapped reads) normalization and log_2_ transform were performed on raw count data.

### Gene fusion detection

STAR-Fusion (v1.4.0) was performed to identify gene fusion events in RNA-seq data^51^. Events with at least one cancer-related gene were included in subsequent analyses. One-sided Wilcoxon rank-sum test was performed to detect fusion events with significantly change of expression in fused samples. Visualization of fusion events was performed by R packages circlize^52^ and chimeraviz^53^.

### Protein extraction and digestion

The organoid lines were collected after washing with PBS buffer thrice and lysed in SDS lysis buffer (4% SDS, 100 mM Tris-HCl, 0.1 M DTT, pH 7.6). The protein was extracted by sonication (JY92-IIDN, Ningblio Scientz Biotechnology Co., LTD, China) at 15% amplitude for 5 s on and 5 s off with the total working time of 2 min. The proteins were then denatured and reduced at 95°C for 5 min. The insoluble debris was removed by centrifugation at 12,000 g for 10 min and the supernatant was retained for proteomic experiment. The protein concentration was determined using fluorescence emission at 350 nm using an excitation wavelength of 295 nm. The measurements were performed in 8 M urea using tryptophan as the standard.

The samples were subjected to in-solution trypsin digestion. Briefly, 1 mg protein was alkylated with iodoacetamide (IAA) in darkness (∼200 μL total volume). The protein was precipitated with 1 mL precooled buffer containing 50% acetone, 50% ethanol, and 0.1% acetic acid and kept in -20°C overnight. The mixture was centrifuged at 18000 g for 40min at 4°C. The protein precipitate was washed with 100% acetone and then 70% ethanol with centrifugation at 18000 g at 4°C for 40min. Following that, 500 μL of 100 mM NH4HCO3 was added into each sample and digested with trypsin (Promega) at a ratio of 1:50 twice for 16h at 37°C. The resulting peptide mixture was acidified with trifluoroacetic acid (TFA) with a final concentration of 1% and then desalted with a C18 cartridge column (Waters, Sep-Pak®Vac 1cc, 50 mg tC18 cartridges) following the manufacturer’s instructions. The amount of the final peptides was determined by NanoDrop One (Thermo Scientific). Finally, ∼400-500 μg peptides can be collected for each sample. For each sample, 1 μg peptides were used for DIA-based proteomic analysis and 300 μg peptides were used to enrich phosphopeptides and then intact glycopeptides in sequence.

### Enrichment of phosphopeptides and glycopeptides

The phosphopeptide enrichment was performed using the High-Select™ Fe-NTA kit (Thermo Scientific, A32992) as described previously.^54^ Briefly, the resins of one spin-column were washed with binding/washing buffer (80% ACN, 0.1% TFA) thrice and divided into 15 aliquots. One aliquot of Fe-NTA was incubated with 300 μg peptides in 300 μL binding buffer for 30 min at room temperature, and then the mixture was transferred into a filter tip (TF-20-L-R-S, Axygen). The supernatant was then removed by centrifugation at 50∼200 g for 1 min. The resins were washed sequentially with 200 μL washing buffer thrice and 200 μL H2O twice to remove the nonspecifically absorbed peptides. The enriched phosphopeptides were eluted from the resins with 200 μL elution buffer (50% ACN, 5% NH3•H2O) twice. The eluted peptides were subjected to vacuum freeze drying and further desalted using C18 StageTips. Half of the purified phosphopeptides were resolved in 0.1% formic acid (FA) and analyzed by LC-MS/MS.

The flow-throughs (FTs) after phosphopeptides enrichment were dried by vacuum freezing and then the glycopeptides were enriched by zwitterionic hydrophilic interaction liquid chromatography (ZIC-HILIC) method. Briefly, the peptides were resuspended in 300 μL loading buffer (80% ACN, 1% TFA) and then loaded onto a homemade Stage-Tip containing 25 mg of ZIC-HILIC particles (Millipore SeQuant® ZIC®-HILIC 3.5 μm, 200 Å) packed onto a C8 disk followed by centrifugation at 2,000 g for 6min. The flow-through peptides were reloaded onto the column for another four times. The Stage-Tips were washed with 200 μL loading buffer for four times. Finally, the enriched glycopeptides were eluted with 140 μL 0.1% TFA and lyophilized for LC-MS/MS. Half of the enriched glycopeptides were resolved in 0.1% formic acid (FA) and analyzed by LC-MS/MS.

### LC-MS/MS analysis

For proteomic analysis, the DIA-based method was used for relative proteome quantification as described previously^55^. For each sample, ∼1 μg peptides were subjected to LC-MS/MS analysis. The peptides were resolved in 0.1% FA and separated using a home-made micro-tip column (75 μm × 200 mm) packed with ReproSil-Pur C18-AQ, C18 3.0 μm resin (Dr. Maisch GmbH, Germany) on a nanoflow HPLC Easy-nLC 1000 system (Thermo Fisher Scientific), using a 90 min LC gradient at 300 nL/min. Buffer A consisted of 0.1% (v/v) FA in H2O and Buffer B consisted of 0.1% (v/v) FA in acetonitrile (ACN). The gradient was set as follows: 3%–27% B in 72 min; 27%–35% B in 10 min; 35%–100% B in 2 min; 100% B in 6 min. Proteomic analyses were performed on a Q Exactive HF mass spectrometer (Thermo Fisher Scientific). The spray voltage was set at 2,300 V in positive ion mode and the ion transfer tube temperature was set at 300°C. Data-independent acquisition was performed using Xcalibur software in profile spectrum data type. The MS1 full scan was set at a resolution of 120,000 @ m/z 200, AGC target 3e6 and maximum IT 100 ms by orbitrap mass analyzer (350-1500 m/z), followed by 40 DIA isolation windows with variable width which were decided by the searching result of a preliminary DDA experiment. The isolation windows were set with 1 Da overlap as follows: 1 loop count of 33 m/z with central m/z at 366; 5 loop counts of 17 m/z with central m/z at 390, 406, 422, 438, 454; 20 loop counts of 13 m/z with central m/z at 468, 480, 492, 504, 516, 528, 540, 552, 564, 576, 588, 600, 612, 624, 636, 648, 660, 672, 684, 696; 8 loop counts of 21 m/z with central m/z at 712, 732, 752, 772, 792, 812, 832, 852; 2 loop counts of 31 m/z with central m/z at 877, 907; 2 loop counts of 51 m/z with central m/z at 947, 997; 1 loop count of 81 m/z with central m/z at 1062; 1 loop count of 401 m/z with central m/z at 1302. The MS2 scans generated by HCD fragmentation at a resolution of 30,000 @ m/z 200, AGC target 5e5 and maximum IT 45 ms. The fixed first mass of MS2 spectrum was set as dynamic. The normalized collision energy (NCE) was set at NCE 27%.

For phosphoproteomic analysis, the DIA-based method was used for relative phosphosites quantification. The LC-MS system was nearly the same as the above proteomic analysis, except the following modifications. The LC gradient was set as follows: 2%–22% B in 72 min; 22%–28% B in 10 min; 28%–100% B in 2 min; 100% B in 6 min. A total of 32 DIA isolation windows with variable width were used for phosphoproteomic analysis. The isolation windows were set with 1 Da overlap as follows: 1 loop count of 57 m/z with central m/z at 378.5; 2 loop counts of 27 m/z with central m/z at 419.5, 445.5; 19 loop counts of 17 m/z with central m/z at 466.5, 482.5, 498.5, 514.5, 530.5, 546.5, 562.5, 578.5, 594.5, 610.5, 626.5, 642.5, 658.5, 674.5, 690.5, 706.5, 722.5, 738.5, 754.5; 4 loop counts of 23 m/z with central m/z at 773.5, 795.5, 817.5, 839.5; 2 loop counts of 33 m/z with central m/z at 866.5, 898.5; 2 loop counts of 47 m/z with central m/z at 937.5, 983.5; 1 loop count of 79 m/z with central m/z at 1045.5; 1 loop count of 417 m/z with central m/z at 1292.5.

For glycoproteomic analysis, the DDA-based label free quantification method was used for the identification and quantification of glycopeptides. The peptides were separated using a home-made C18 3.0 μm micro-tip column on a nanoflow HPLC Easy-nLC 1200 system (Thermo Fisher Scientific) with a 120 min LC gradient at 300 nL/min. Buffer A consisted of 0.1% (v/v) FA in H2O and Buffer B consisted of 0.1% (v/v) FA in 80% acetonitrile (ACN). The gradient was set as follows: 2%-5% B in 1 min; 5%–32% B in 94 min; 32%–45% B in 15 min; 45%–100% B in 2 min; 100% B in 8 min. Proteomic analyses were performed on a Q Exactive HF-X mass spectrometer (Thermo Fisher Scientific). The spray voltage was set at 1,800 V in positive ion mode and the ion transfer tube temperature was set at 320°C. Data-independent acquisition was performed using Xcalibur software in profile spectrum data type. The MS1 full scan was set at a resolution of 60,000 @ m/z 200, AGC target 3e6 and maximum IT 20 ms by orbitrap mass analyzer (300-1800 m/z), followed by ‘top 20’ MS2 scans generated by HCD fragmentation at a resolution of 15,000 @ m/z 200, AGC target 1e5 and maximum IT 100 ms. The fixed first mass of MS2 spectrum was set as dynamic. Isolation window was set at 1.4 m/z. The stepped normalized collision energy (NCE) was set at 20%, 30%, 40%, and the intensity threshold was set to 2.0e4. The dynamic exclusion time was 45 s. Precursors with charge 1, 7, 8 and > 8 were excluded for MS2 analysis.

### Database searching of MS data

The “DirectDIA” pipeline in Spectronaut v15 software was used to analyze the proteomic and phosphoproteomic data in a spectral library-free mode with the default parameters. All DIA runs were directly searched against the Swiss-Prot protein database (downloaded in June, 2021, 20,600 entries). For proteomic analysis, Trypsin/P was set as the enzyme and peptide length from 7-52. Carbamidomethyl (C) was set as the fixed modification, and acetyl (protein N-term) and oxidation (M) was set as the variable modifications. The max variable modifications were set to 5. The FDRs of PSM, peptide and protein group were all set to 1%. The precursors were filtered by Qvalue. For phosphoproteomic analysis, and phospho(S/T/Y) was set as another variable modification and the PTM localization option was enabled. The PTM localization score >0.75 was used as the criteria of highly reliable Class-I phosphosites.

The glycoproteomic data was analyzed and quantified by label free quantification using the PEAKS GlycanFinder software. Only glycopeptides with a glycosylation site specific accuracy A-Score >20 were retained for the following analysis, resulting in 38,691 glycopeptides.

### Data quality control

We combined dozens of organoid lines to get a mixed sample, which was separated into 1 mg aliquotes and used as quality control samples. The organoid protein samples were subjected to trypsin digestion in several batches with one control sample in every batch. The repeatability in the proteomic and phosphoproteomic quantification of the mixed samples was used to describe the data quality.

### Proteogenomic data processing

Proteins and phosphosites were log_2_ transform and were median normalized on raw abundance profile. Proteins and phosphosites missing in more 50% samples were filtered out. Perseus software (v1.6.15.0) was used to do missing data imputation based on normal distribution. After filtering, we obtained normalized abundance profile of 7225 proteins and 17781 phosphosites. Combat function in SVA package was used to correct the batch effects on proteomics and phosphoproteomics data^56^. In addition, the phosphoproteomic profile was normalized according to the abundance of corresponding proteins.

### Glycoproteomic Data Processing

We have detected 38691 glycopeptides and we categorized these glycopeptides into four glycan types: high-Mann (oligomannose), fucose (fucosylated glycans), sialic (sialylated glycans; including glycans with sialylation only and those with both sialylation and fucosylation) and others (other glycoforms) according to published research^57^. Log_2_ transform and median normalization were then performed on raw abundance profile. Glycopeptides missing in more 30% samples were filtered out. Perseus software (v1.6.15.0) was used to do missing data imputation based on normal distribution. After filtering, we obtained normalized abundance profile of 2662 glycopeptides.

To compare proteoglycan level among different groups, ANOVA/t-test was performed to identify differential glycopeptides among different subtypes. At the glycosylation site level, mean abundance of each differential glycopeptides in each group is calculated and paired t-test was then performed to measure the difference. At the sample level, average of all differential glycopeptides in each sample is calculated and t-test was then performed to measure the difference.

### Proteogenomic-based subtyping

Unsupervised subtyping was conducted with consensus clustering by R package “CancerSubtypes” (function: ExecuteCC; clusterAlg=“km”; reps=10000) on normalized abundance profile of top 50 percent variance proteins (n=3612) of 93 PDAC samples.^58^ Differential proteins among proteomic subtypes were performed by ANOVA (FDR<0.05; FC≥1.5). Gene set enrichment analysis (GSEA) was performed on ranked proteins (ranked based on signed p-value of the Student’s t test) to determine pathways enriched in different proteomic subtypes (R package “clusterProfiler”, FDR<0.1)^59,60^. Molecular Signatures Database (MSigDB) of KEGG pathways (“c2.cp.kegg.v7.5.symbols.gmt”) were used for enrichment analysis^61^. Proteoglycan pathway related genes were defined as genes enriched in GSEA enriched glycan-related pathways in Glycan-H subtype (Glycosaminoglycan Degradation, N-Glycan Biosynthesis, Glycosaminoglycan Biosynthesis Keratan Sulfate, Glycosylphosphatidylinositol Gpi Anchor Biosynthesis, O-Glycan Biosynthesis and Glycosaminoglycan Biosynthesis Heparan Sulfate). In addition, gene set variation analysis (GSVA) was also processed to calculate GSVA enrichment scores of KEGG pathways for each sample on both protein and mRNA profile^62^. Then spearman correlation analysis was performed on GSVA scores across PDAC samples to evaluate consistency between transcriptomic and proteomic on pathway level.

### Validation of proteomic subtypes

To further validate the three proteomic subtypes, we reanalyzed two published PDAC cohorts: the Hyeon cohort^15^ and the Cao cohort^16^. Using the same classification method (Consensus clustering using R package “CancerSubtypes”, function: ExecuteCC; clusterAlg=“km”; reps=10000), we identified three protein subtypes in the Hyeon cohort. Through NearestTemplatePrediction (NTP)^63^, the Cao cohort was stratified into three subtypes based on differential proteins among three proteomic subtypes in our cohort. Only 105 samples with sufficient tumor cellularity were included. GSVA enrichment scores for KEGG pathways were calculated on these two cohorts GSVA enrichment scores for KEGG pathways were calculated on these two cohorts, and differentially enriched pathways were identified by ANOVA with FDR < 0.01^62^.

### Drug screening

Organoids were digested with Tryp-LE, and inactivated digestive enzyme with 1640 basic medium with 10% fetal bovine serum (FBS). Before being resuspended in medium, the cells were washed with cold PBS twice. Organoids were transferred into 384-well plates in 50μl of medium using Multidrop (Thermo) dispensers (3000 cells per well). Then, we added the drugs with 7 different concentration gradients (10-fold dilutions series or 8-fold dilutions series) in the next day by mosquito workstation (SPT LabTech). Cell viability was assayed by Envision (Perkinelmer) using CellCounting-Lite® 2.0 (Vazyme) after five days treatment. Effects on cell viability were normalized according to DMSO. Each drug concentration (umol/L) was log_10_ transformed. Average inhibition rates from two independent experiments were calculated with Excel and visualized using GraphPad Prism 9. The AUC and IC50 was calculated with the GraphPad Prism 9. The normalized AUC was obtained by dividing one AUC by the maximum AUC for each drug. IC50 was normalized by log_2_ transformation. Spearman correlation analysis was performed on IC50 and AUC profile across samples to measure consistency between IC50 and AUC. In addition, Spearman correlation analysis was also performed on biological replicates to measure the reproducibility of independent screens.

### Classifying synergy

To detect synergy, we compared observed combination responses to expected combination responses. For the latter, we used Bliss independence of the response to the statins and the chemos alone. Conceptually, every point on the Bliss dose response curve is defined as the product between the statins viability and the corresponding point on the chemos dose response curve^64^. Shifts in potency (ΔAUC) were calculated as the difference between the observed combination response and expected Bliss (ΔAUC = Bliss AUC − combination AUC). ΔAUC > 0 represented synergistic.

### Chemotherapeutic based subgrouping

By unsupervised hierarchical clustering (clustering method= “ward.D2”), drug response data (measured by AUC) of 5 chemotherapeutic agents (5-FU, GEM, IRI, OXA and PTX, commonly used to treat PDAC patients) were clustered into three groups: resistant group, intermediate group and sensitive group. GSEA was performed to identify pathway-level features of chemoresistant and chemosensitive groups.

### Prediction of patient treatment response

In order to predict patient treatment responses, we have developed organoid drug sensitivity scoring criteria. Firstly, based on the results of organoid drug sensitivity tests, individual drugs in the chemotherapy regimen are assigned sensitivity scores. “Resistant” is scored as 0, “median” as 1, and “sensitive” as 2. The sensitivity score for the chemotherapy regimen is then calculated as the sum of the scores for the individual drugs. If the total sensitivity score for the chemotherapy regimen is ≥ 2, it predicts a patient response to the regimen; otherwise, it predicts no response (refer to Table 1).

The assessment of the actual treatment responses in patients is based on imaging examinations during treatment. Using the RECIST v1.1 criteria^65^, patients’ treatment responses are classified as partial responses (PR), stable disease (SD), and progressive disease (PD). PR and SD are considered responsive to treatment. By comparing the predicted results with the actual treatment responses, the accuracy of the organoid drug sensitivity score in predicting treatment responses can be evaluated.

### Immunohistochemistry and immunofluorescence

Immunohistochemistry and immunofluorescence were performed as described in our previous study^66^. The antibodies used for staining was as follows: Anti-SNA (1:400, vector laboratories, B-1305-2), anti-Ecad (1:400, CST, 14472S).

### Detection of cholesterol levels in organoids

The cultured organoids were harvested and washed once with ice-cold PBS to remove residual media. Cell counting was performed using an automated cell counter, and 1,000,000 cells were aliquoted for subsequent analyses. The cholesterol content in the cells was quantified using the Amplex Red Cholesterol and Cholesteryl Ester Assay Kit (Beyotime Biotechnology, cat# S0211S), following the manufacturer’s instructions.

### qRT-PCR

qRT-PCR was performed as described in our previous study.^67^ The primers used for each of the genes are listed (Supplementary Table 5).

## Data Availability Statements

All raw sequencing data has been deposited in Genome Sequence Archive (GSA, accession number HRA007290) database and National Omics Data Encyclopedia (NODE, accession number OEP004966) database. All data is publicly accessible. Proteomic, phosphoproteomic and glycoproteomic data in this paper have been deposited in the OMIX (https://ngdc.cncb.ac.cn/omix: accession no. OMIX006354, no. OMIX006355, no. OMIX006356).

## Code Availability Statements

This paper does not report original code. Any additional information required to reanalyze the data reported in this work paper is available from the lead contact upon request.

## Notes

https://ngdc.cncb.ac.cn/gsa/

https://ngdc.cncb.ac.cn/omix

### Competing Interest Statement

The authors have declared no competing interest.

