## Extended figures for "Targeting protein glycosylation and cholesterol metabolism in chemoresistant pancreatic cancer"

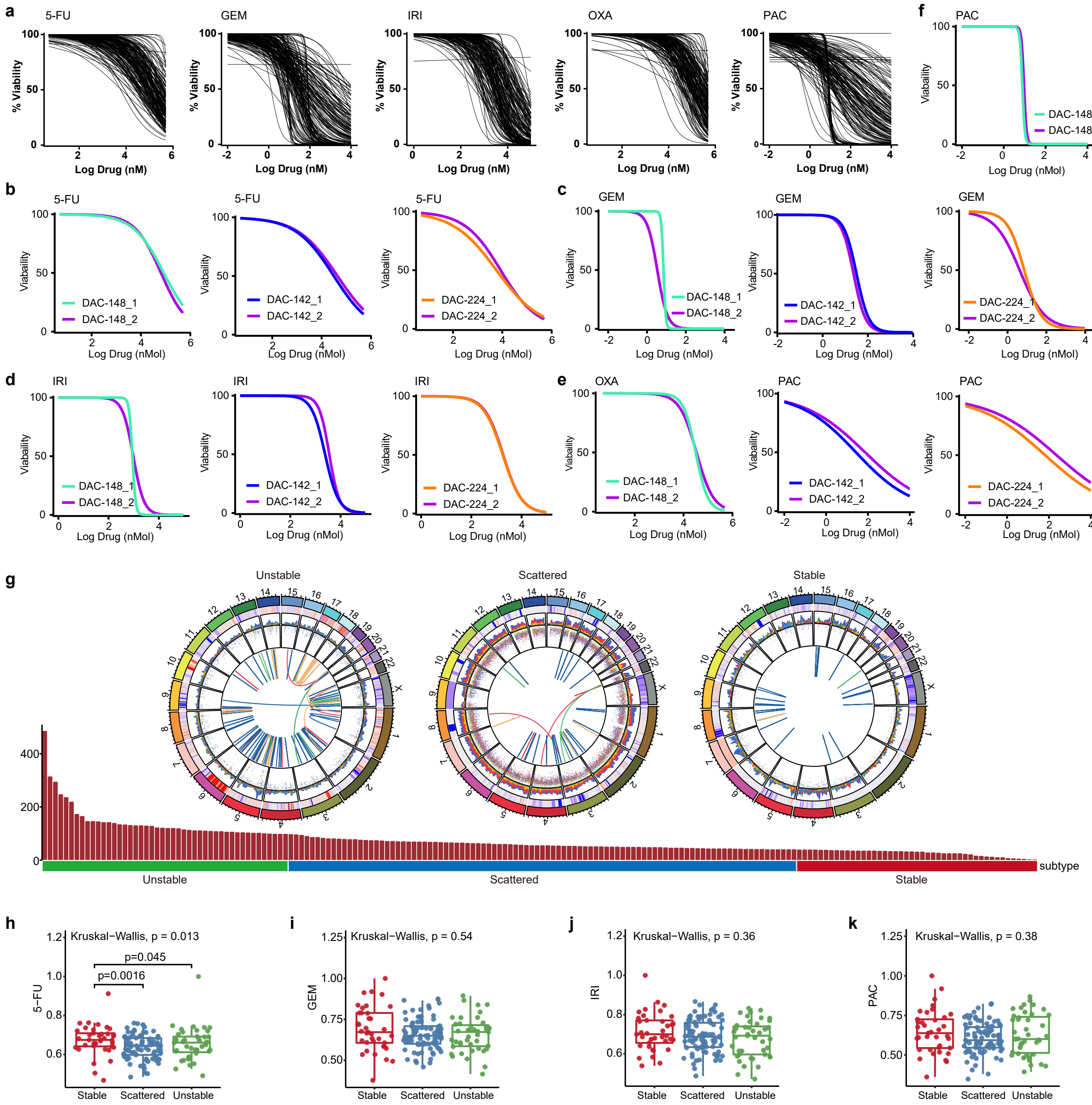


**Extended Data Fig. 1. Therapeutic profiling of PDAC organoids using the 5 chemotherapeutic agents. a**, Dose-response curves for 5-FU, GEM, IRI, OXA and PAC. **b-f**, Dose-response curves of 5-FU (**b**), GEM (**c**), IRI (**d**), OXA (**e**) and PAC (**f**) was stable over multiple passages in the indicated organoid lines. **g**, Bar plot showing structural variants numbers for ‘stable’, ‘scattered’ and ‘unstable’ subtypes (bottom) and representative tumors of each group (top). The colored outer rings represent chromosomes. The subsequent ring illustrates copy number variations, with red indicating gains and blue indicating losses. The following ring displays the inter-variant distances at genomic loci (display hyper mutated genomic regions by rainfall plot). The innermost lines depict chromosomal structural rearrangements. **h-k**, Normalized AUC of 5-FU (**h**), GEM (**i**), IRI (**j**) and PAC (**k**) among ‘stable’, ‘scattered’ and ‘unstable’ subtypes, P-values are calculated by Kruskal-Wallis test.

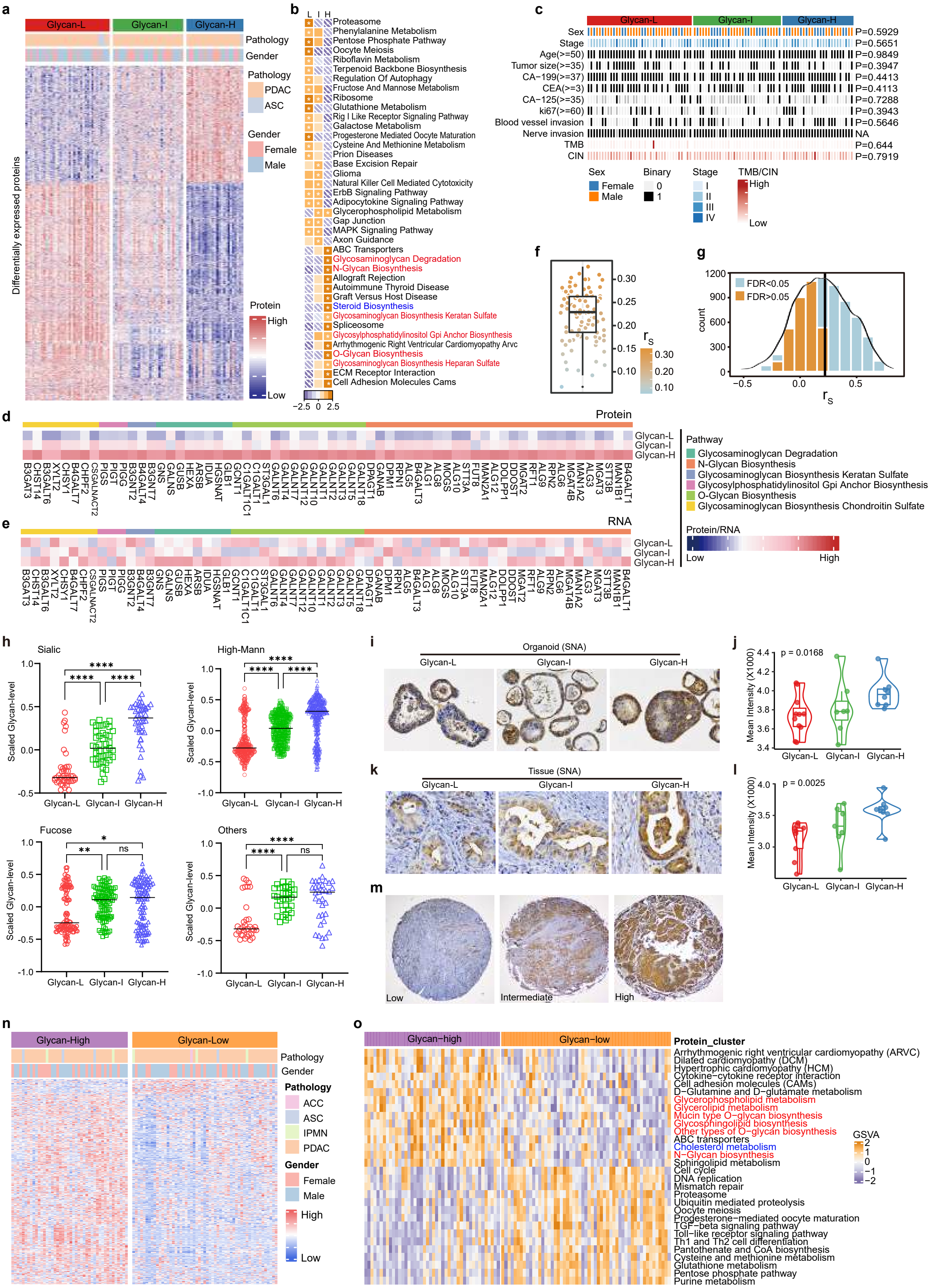


**Extended Data Fig. 2. Proteomic subtypes with strong prognostic and chemosensitivity implications. a,** Heatmap showing differential proteins among three proteomic subtypes (ANOVA, FDR < 0.05; FC ≥ 1.5). **b,** Pathway enrichment analysis among three proteomic subtypes. Shown are normalized enrichment score (NES) scores of KEGG pathways calculated by GSEA based on ranked proteins (ranked based on signed p-value of t test, one compared to others). Asterisks indicate gene sets with $FDR < 0.1$. **c**, The distribution of clinical parameters for each patient of three proteomic subtypes. P values are calculated by fisher's exact test. **d**, Heatmap showing mean protein abundance of glycan pathway genes (enriched in Glycan-H by GSEA) among three proteomic subtypes. **e**, Heatmap showing mean RNA expression of glycan pathway genes among three proteomic subtypes. **f**, Boxplot of RNA-to-protein correlations in each PDAC sample calculated by Spearman correlation analysis. Scale bar shows the Spearman's rho. In the box plots, the central line represents median; bounds of box represent the first and third quartiles, and the upper and lower whiskers extend to the highest or the smallest value within 1.5 times interquartile range (IQR); P values are adjusted by Benjamini-Hochberg method. **g**, Histogram and density distribution of gene-wise RNA-to-protein correlations calculated by Spearman correlation analysis. P values are adjusted by Benjamini-Hochberg method. FDR of correlation test less than 0.05 shown in blue, and more than 0.05 shown in orange. **h**, Comparison of glycan level (Z-score) of four types of glycosylated proteins (Sialic, High-Mann, Fucose and Others) among three proteomic subtypes at the glycosylation site level, differential glycopeptides with $p < 0.05$ were included in the analysis. P-values are calculated by t-test (*: $p < 0.05$; **: $p < 0.01$; ***: $p < 0.001$; ****: $p < 0.0001$). **i**, Representative images of immunohistochemical staining show low, intermedium and high sialic acid (SNA) staining in organoid lines. **j**, Mean intensity of SNA among three proteomic subtypes in organoid lines. P-value is calculated by t-test. **k**, Representative images of immunohistochemical staining show low, intermedium and high SNA staining in PDAC tissues. **l**, Mean Intensity of SNA among three proteomic subtypes in tissue. P-value is calculated by t-test. **m**, Representative images of immunohistochemical staining show low, intermedium and high SNA staining in Tissue Microarray (TMA). **n**, Heatmap showing differential glycopeptides among two glycoproteomic subtypes (t-test, $p < 0.05$). **o**, Heatmap showing pathways with differential GSVA enrichment scores among two glycoproteomic subtypes (t-test, $FDR<0.01$).

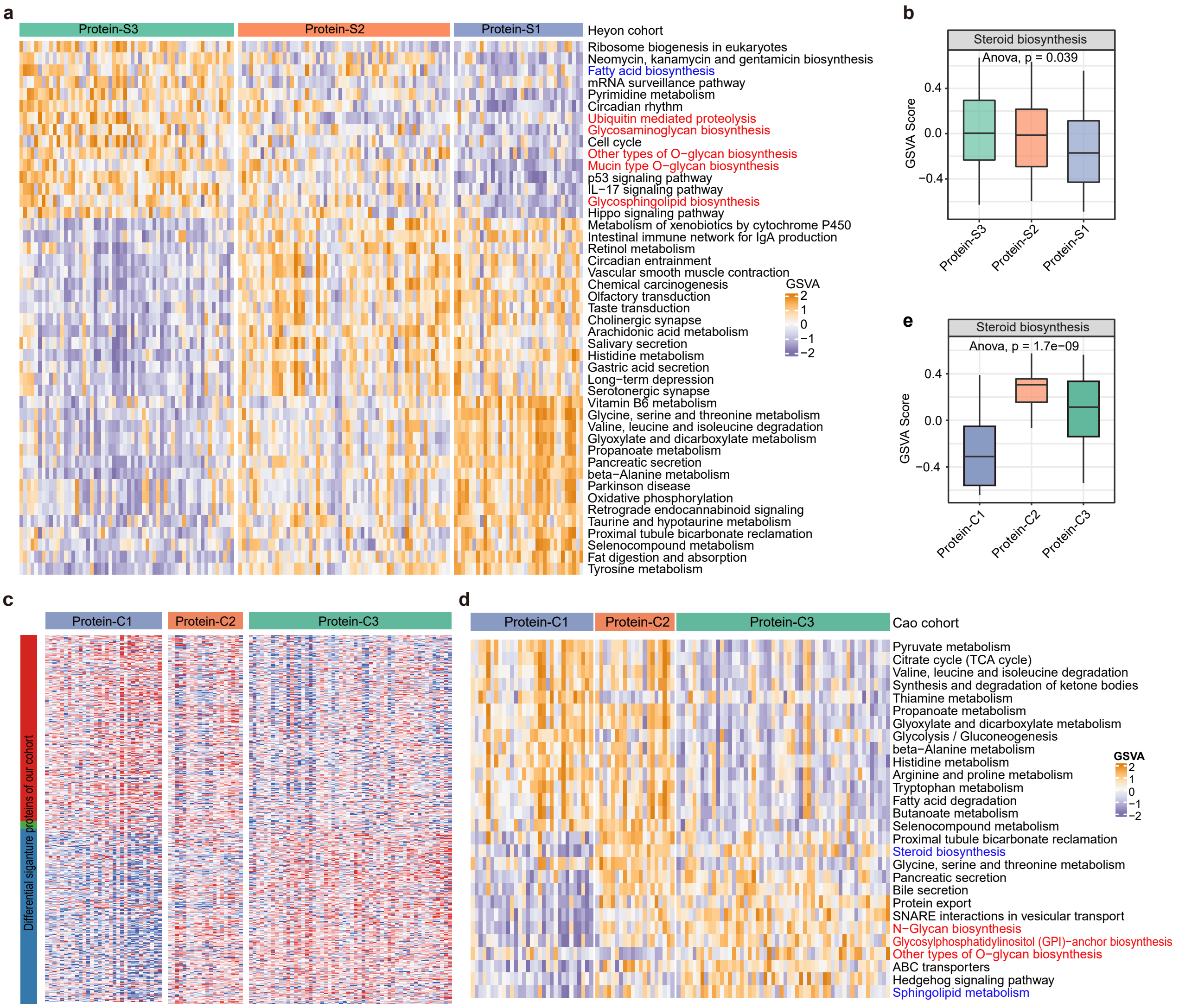


**Extended Data Fig. 3. Validation of proteomic subtypes. a**, Heatmap showing pathways with differential GSVA enrichment scores among three proteomic subtypes on Hyeon cohort (ANOVA, FDR<0.01). **b**, Boxplot showing GSVA enrichment scores for steroid biosynthesis pathway among three proteomic subtypes on Hyeon cohort. **c**, Cao cohort were dividing into three subgroups based on differential proteins among three proteomic subtypes of organoids. **d**, Heatmap showing pathways with differential GSVA enrichment scores among three proteomic subtypes on Cao cohort (ANOVA, FDR<0.01). **e**, Boxplot showing GSVA enrichment scores for steroid biosynthesis pathway among three proteomic subtypes on Cao cohort.

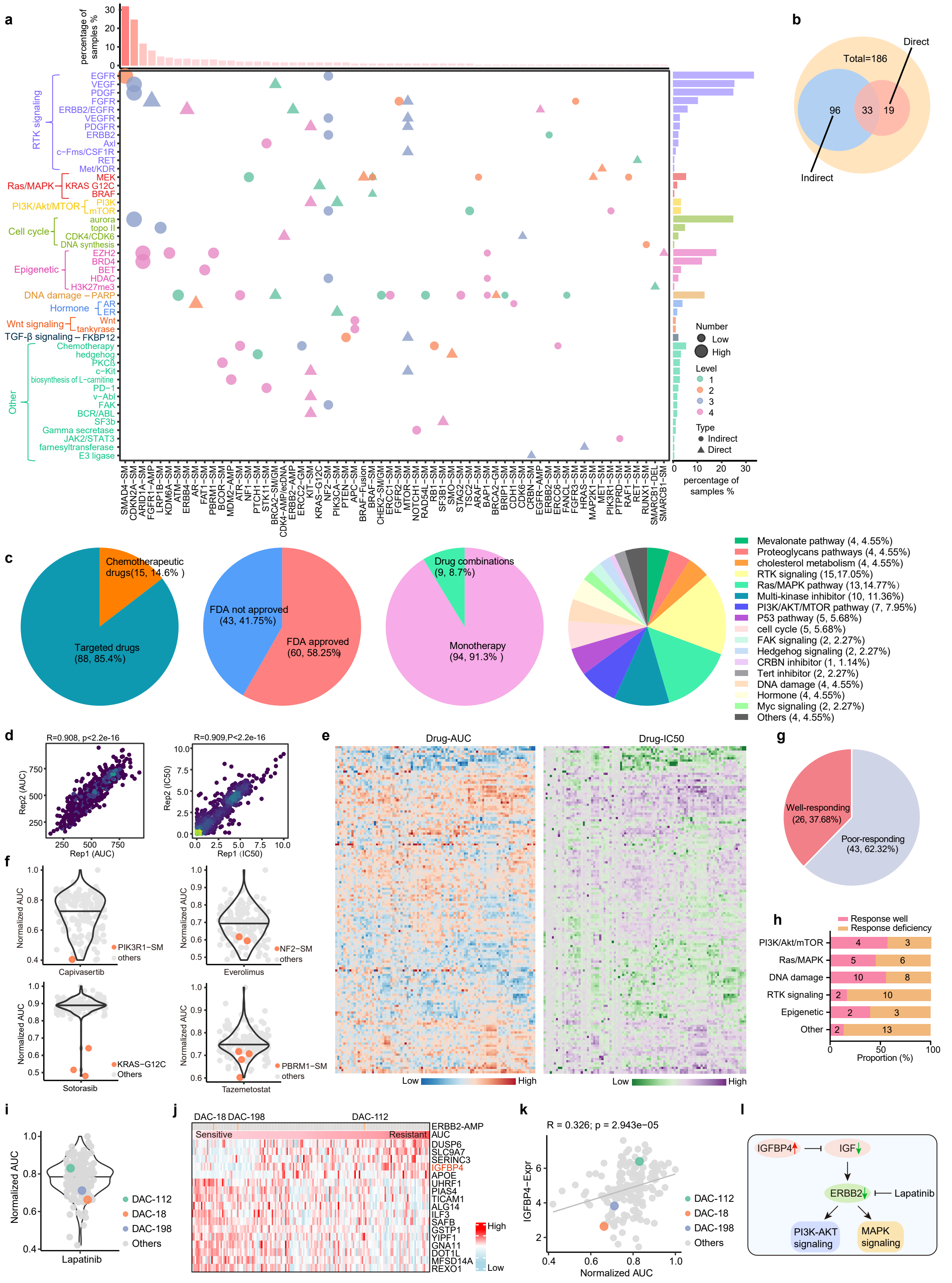


**Extended Data Fig. 4. A landscape of therapeutic strategies for PDAC through pharmacogenomic insights. a,**

Relationship between AGVs and drug targets (classified based on targeted pathways) according to OncoKB and CGI evidence tiers for pan-cancer. **b**, Pie chart showing proportion of PDAC patients with clinically AGVs (direct or indirect). **c**, Pie charts showing classification of drugs for therapeutic profiling in our cohort. **d**, Spearman correlation plot between biological replicates of 10 organoid lines from independent screens. **e**, Heatmap showing matched AUC (left) and IC50 profile (right). Values are normalized by Z-score and each column represents a drug while each row represents a sample. **f**, Violin plot showing normalized AUC of Capivasertib, Everolimus, AMG510 and Tazemetostat. Samples with corresponding AGVs are marked in orange. **g**, Pie chart showing proportions of well-responding AGV-drug pairs and poor-responding AGV-drug pairs. **h**, Proportions of well-responding AGV-drug pairs and poor-responding AGV-drug pairs based on pathways. **i**, Violin plot showing normalized AUC of Lapatinib and three samples with *ERBB2* amplification are marked. **j**, RNA expression of genes highly correlated with normalized AUC of Lapatinib performed by spearman correlation analysis ($R \geq 0.3$ and $p < 5e^{-5}$). **k**, Correlation plot between expression of *IGFBP4* and normalized AUC of Lapatinib performed by spearman correlation analysis. **l**, Schematic illustrating how *IGFBP4* overexpression induces lapatinib resistance in organoids.

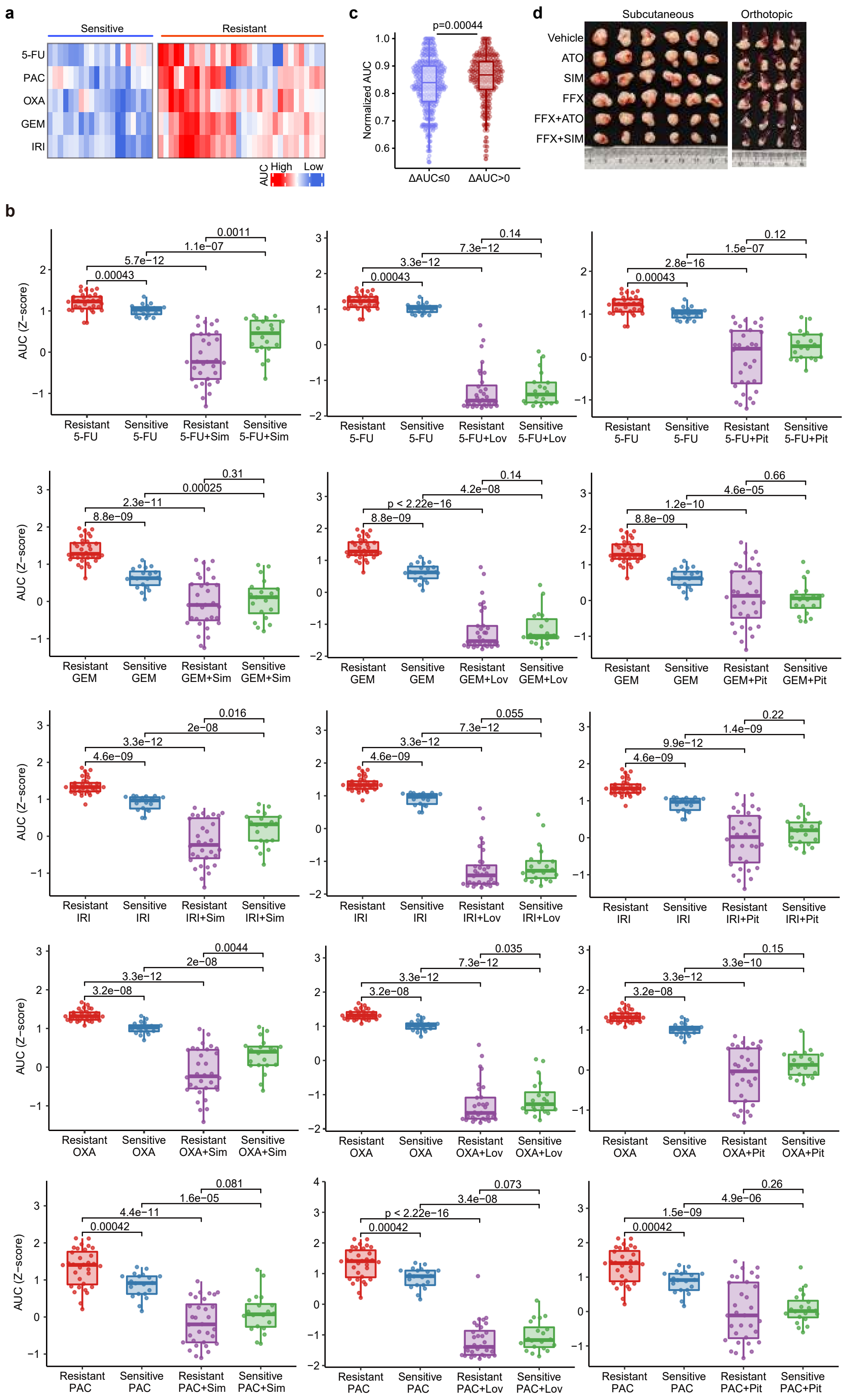


**Extended Data Fig. 5. Statins overcome chemoresistance in PDAC. a**, Heatmap showing AUC profile of 5 chemotherapeutic drugs which are clustered into two subtypes in validation set. **b**, AUC (Z-score) comparison for chemotherapy only and combination of chemotherapy and statins among the two chemotherapeutic subtypes in validation set for each drug pair. P-value is calculated by one-sided Wilcoxon rank-sum test. **c**, Distribution of normalized AUC (chemotherapeutic drugs) of synergistic and non-synergistic organoids in validation set. P-value is calculated by one-sided Wilcoxon rank-sum test. **d**, Image of FFX resistant organoid (DAC-71) derived subcutaneous xenografts (left) and orthotopic xenograft (right) isolated from each group with indicated drug treatment.

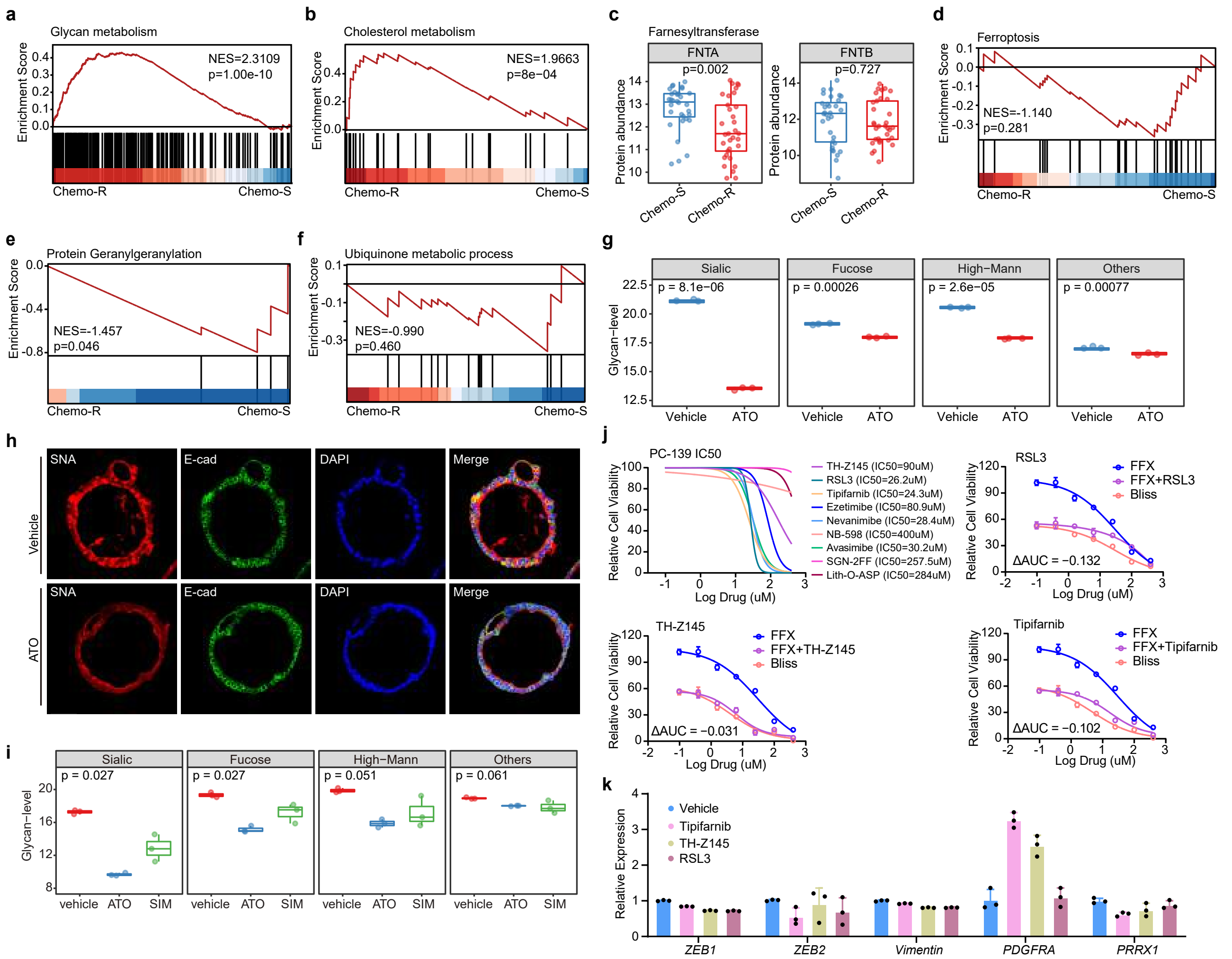


**Extended Data Fig. 6. The mechanism of statins enhance cellular sensitivity to chemotherapeutic agents. a**, GSEA enrichment plot for a gene set comprising all KEGG glycan-related pathways in Chemo-R group versus Chemo-S group. **b**, GSEA enrichment plot for Cholesterol metabolism pathway in Chemo-R group versus Chemo-S group. **c**, Boxplot of Farnesyltransferase FNTA and FNTB expression levels stratified by chemotherapeutic subtypes. P-values are calculated by t-test. **d**, GSEA enrichment plot of ferroptosis in Chemo-R group versus Chemo-S group. **e**, GSEA enrichment plot of protein geranylgeranylation in Chemo-R group versus Chemo-S group. **f**, GSEA enrichment plot of ubiquinone metabolic process in Chemo-R group versus Chemo-S group. **g**, Comparison of protein glycosylation levels of four types of glycosylated proteins (Sialic, High-Mann, Fucose and Others) among ATO treated and control organoids. P-values are calculated by t-test. **h**, Immunofluorescence staining for SNA and E-cadherin (E-cad) shows that ATO inhibit protein glycosylation of the indicated organoid line (DAC-71). Scale bar = 50 μm. **i**, Comparison of protein glycosylation levels of four types of glycosylated proteins (Sialic, High-Mann, Fucose and Others) among ATO treated, SIM treated and control group for the DAC-71 derived subcutaneous xenografts. P-values are calculated by ANOVA. **j**, Dose response curves of FFX in combination with inhibitors targeting other pathways downstream of the MVA in the Chemo-R group organoid line DAC-36. **k**, Bar plot showing the changes in expression levels for EMT signature genes after treatment with inhibitors targeting ferroptosis, farnesylated proteins and protein prenylation pathways downstream of the MVA in the DAC-36 organoid line.

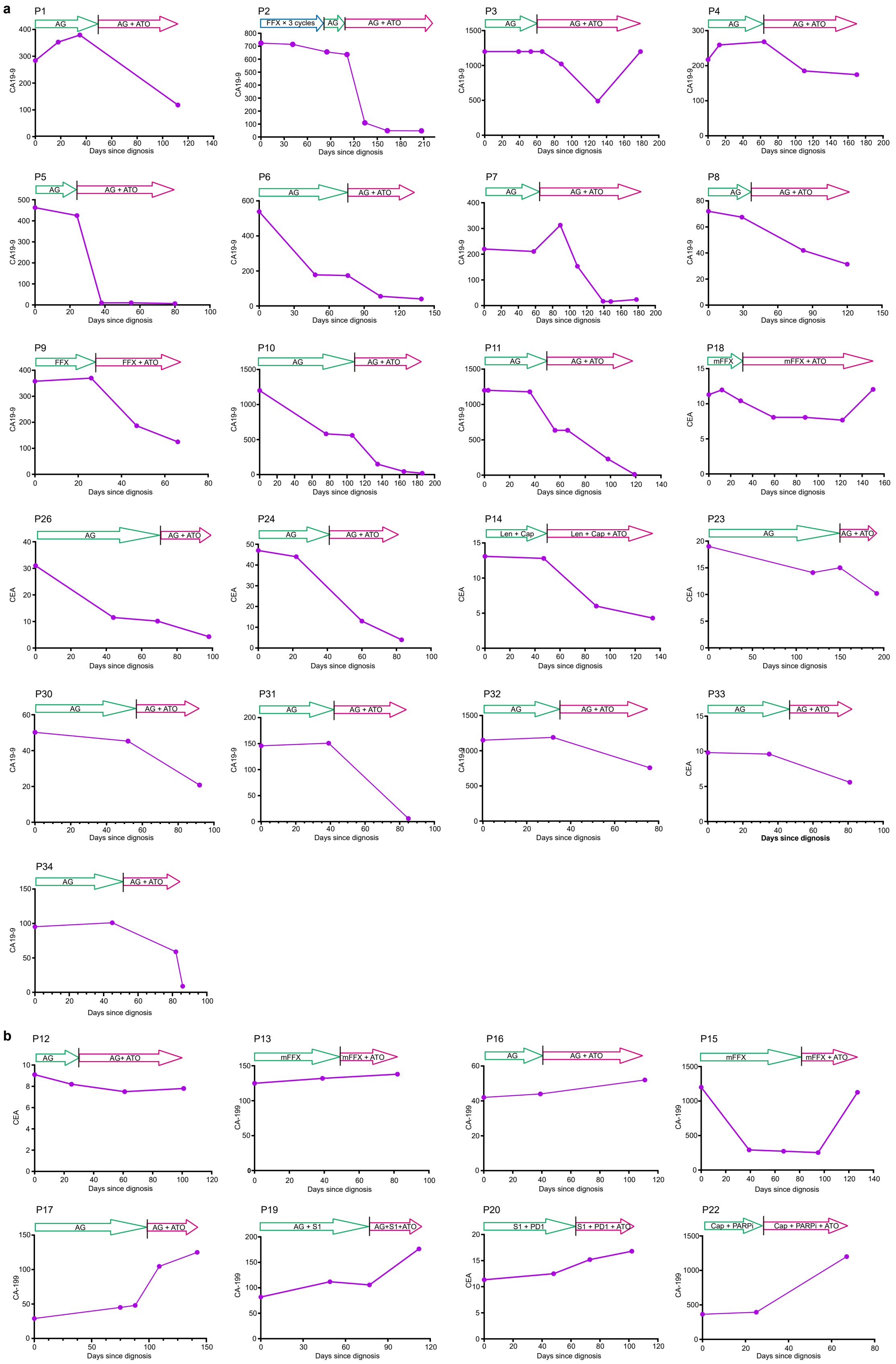


**Extended Data Fig. 7. Statins improve chemotherapy efficacy in clinical treatment. a-b**, Serum CA19-9 or CEA measurements from diagnosis throughout the patient's treatment course. The arrow indicates the corresponding treatment method. Patients with CA19-9 or CEA decrease > 20% are shown in (**a**), while patients whose CA19-9/CEA remained stable or increased are shown in (**b**).
